# Decoding and perturbing decision states in real time

**DOI:** 10.1101/681783

**Authors:** Diogo Peixoto, Jessica R. Verhein, Roozbeh Kiani, Jonathan C. Kao, Paul Nuyujukian, Chandramouli Chandrasekaran, Julian Brown, Sania Fong, Stephen I. Ryu, Krishna V. Shenoy, William T. Newsome

## Abstract

In dynamic environments, subjects often integrate multiple samples of a signal and combine them to reach a categorical judgment. The process of deliberation on the evidence can be described by a time-varying decision variable (DV), decoded from neural activity, that predicts a subject’s decision at the end of a trial. However, within trials, large moment-to-moment fluctuations of the DV are observed. The behavioral significance of these fluctuations and their role in the decision process remain unclear. Here we show that within-trial DV fluctuations decoded in real time from motor cortex are tightly linked to choice behavior, and that robust changes in DV sign have the statistical regularities expected from behavioral studies of changes-of-mind. Furthermore, we find single-trial evidence for absorbing decision bounds. As the DV builds up, heavily favoring one or the other choice, moment-to-moment variability in the DV is reduced, and both neural DV and behavioral decisions become resistant to additional pulses of sensory evidence as predicted by diffusion-to-bound and attractor models of the decision process.

---

When making a categorical decision about a noisy stimulus, it is common to fluctuate between levels of commitment to a choice before reporting a decision. In some instances the fluctuations are sufficiently strong to lead to a “change of mind” (CoM) while deliberating^1-6^ or even while the reporting action is being executed^7^. Because these within-trial fluctuations are different from trial to trial and not necessarily tied to an external event or stimulus feature, they can only be captured using a moment-to-moment neural readout of the decision state on single trials.

To obtain this readout, we decoded a decision variable (DV) from neural population activity in PMd and M1 in real time to continuously estimate the decision state while two monkeys performed a motion discrimination task^8, 9^ (Fig. 1a, see Methods). The DV was estimated by applying a linear decoder, trained on data from a previous experimental session, to spiking data (from 96 to 192 electrodes) from the preceding 50 ms, updated every 10 ms throughout each trial (Fig. 1b, see Methods). The sign of the DV indicated which choice was predicted by the decoder, which allowed us to calculate the decoder’s prediction accuracy. The DV magnitude reflected the confidence of the model’s prediction in units of log-odds for one vs. the other decision (see Methods). Note that the decision variable as defined here encompasses all choice predictive signals that can be decoded from neural activity^10^, including but not limited to moment-to-moment value of accumulated evidence as posited in classical sequential sampling models.

**Figure 1.**
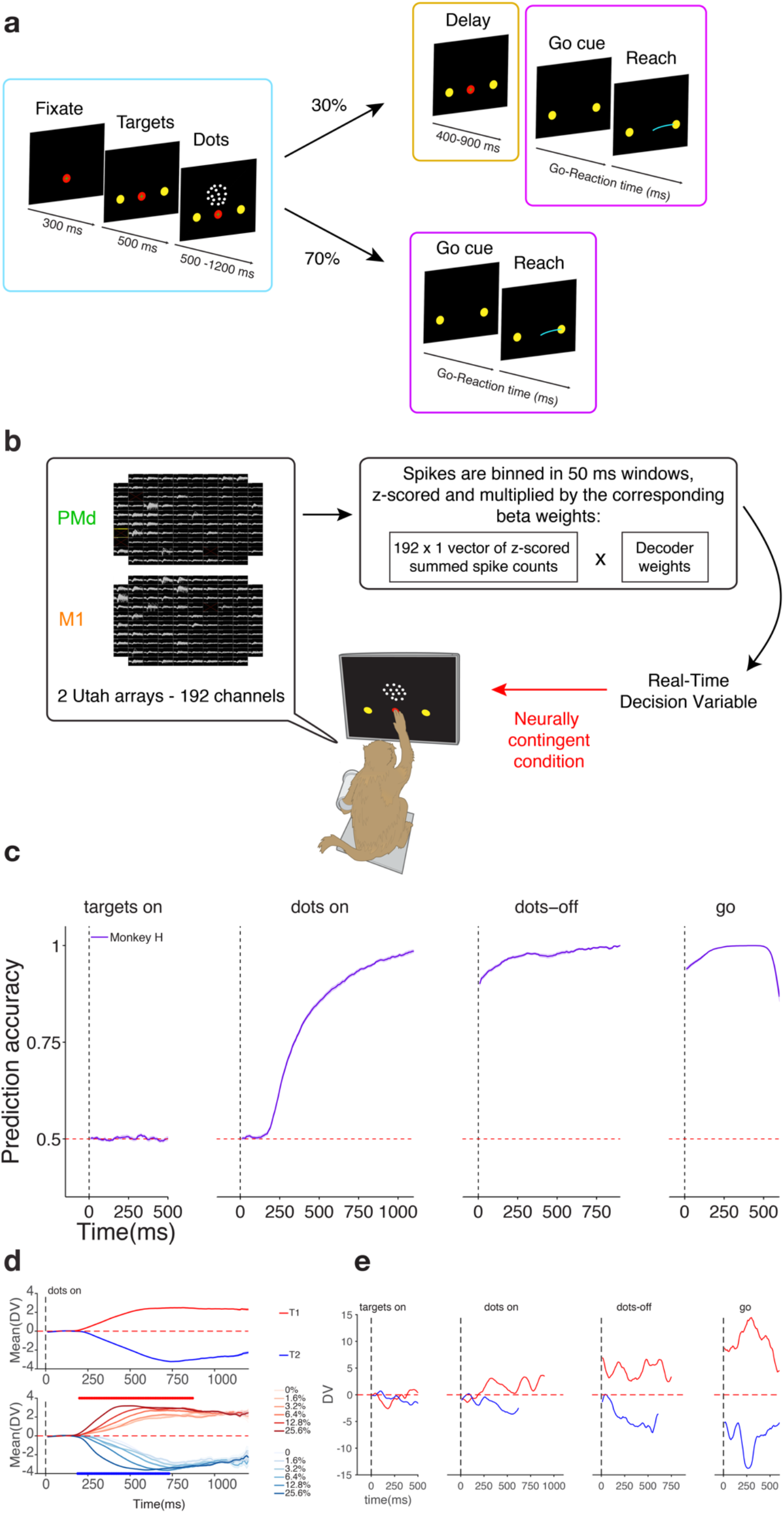
Setup and performance of real-time readout of decision states during a motion discrimination task. **a) Motion discrimination task.** Trials began with the onset of a fixation point (FP) on the touchscreen. Once both eye and hand fixation were acquired, two targets appeared on the screen. The motion stimulus was shown after a short delay (500 ms) and lasted 500-1200 ms for the open-loop trials. On 70% of the trials the dots offset was followed by the go cue (no delay period), while on the remaining 30% the subject was required to withhold a response for a random delay duration (400-900 ms). Decision states were continuously decoded during all epochs of the trial. Three different decoders were used during different trial epochs, shown by the different colored boxes (blue, yellow and purple; see Methods). **b) Real-time, closed-loop setup.** Neural activity from 96-channel Utah Arrays was continuously recorded and processed while monkeys performed the motion discrimination task. For monkey H, two Utah arrays implanted in PMd and M1 were used. For monkey F only one Utah array implanted in PMd was utilized. During data collection, the recorded neural activity was binned, summed, z-scored and projected onto a single dimension: a linear choice decoder. The result of this operation was our real time read out of commitment, which could be used to stop the stimulus presentation in a neurally contingent manner (red arrow), thereby closing the loop in the experiment. **c) Choice prediction accuracy obtained from real-time, open-loop readout.** Average prediction accuracy (see Methods) over time ± SEM for monkey H is plotted in purple. Prediction accuracy is calculated for each time point aligned to four different events in the trial (targets onset, dots onset, dots offset and go cue) using the real-time DV and quantified as the fraction of trials in which the classifier correctly predicted the monkey’s upcoming choice. For logistic regression this operation is equivalent to comparing the DV sign to the choice sign. Accuracy was calculated for each session and averaged across sessions using a total of 16468 trials for monkey H. **d) Average Decision Variable traces during dots period.** Top panel: Average DV during the dots epoch for right (red) and left (blue) choices for monkey H. Bottom panel: Average DV sorted by choice and stimulus coherence (correct trials only) for monkey H. Darker shades correspond to higher stimulus coherence. Red and blue dots indicate timepoints for which coherence was significant regressor of DV for T1 and T2 choices respectively (correct trials only, p<10^-5^ uncorrected). For monkey H coherence is a significant regressor of DV for at least one of the choices for the period between [190, 870] ms aligned to dots onset. **e) Example DV traces captured during open loop trials.** DV traces for two trials are plotted as a function of time aligned to four different events: targets onset, dots onset, dots offset and go cue. The trial in red led to a right choice whereas the trial in blue led to a left choice. Despite the stability in DV sign for these two trials from ∼250 ms after dots onset until the end of the trial, strong fluctuations in DV magnitude were observed in both cases, within and across epochs.

We have previously demonstrated with offline analysis that this decision variable (DV) can predict choices on single trials up to seconds before initiation of the operant response, and that the accuracy of these predictions increases on average throughout the course of the trial^10^.

Here, we employed closed-loop, neurally-contingent control over stimulus timing to directly probe the relationship of within-trial DV fluctuations to behaviorally meaningful decision states. For the first time, we quantified the behavioral effects of previously covert DV variations (i) as a function of time and for different virtual DV boundaries imposed during the trial, (ii) when large, CoM-like fluctuations were detected during deliberation on noisy visual evidence, and (iii) when subthreshold stimulus pulses were added during the trial.

Having a nearly instantaneous real-time estimate of the decision state read-out enabled us to terminate the visual stimulus based on the current value (or history) of the DV and validate the behavioral relevance of DV fluctuations using the monkey’s behavioral reports following stimulus termination.

## Decisions on perceived stimulus motion can be reliably decoded in real time based on 50 ms of PMd/M1 neural activity

Two monkeys performed a variable duration variant of the classical random dot motion discrimination task using an arm movement as the operant response^10^. As expected, the subjects performed better for higher coherence and longer duration stimuli and reached almost perfect performance for the easiest stimuli (Extended Data Fig. 1).

We first measured the accuracy of our real-time decoder in predicting the monkeys’ behavioral choices as a function of time during the trial. As in our previous offline results^10^, average prediction accuracy started at chance levels during the targets epoch (Fig. 1c, Extended Data Fig. 2a). During the dots presentation average prediction accuracy quickly departed from baseline (174.5 ms ± 18.8 and 214.5 ms ± 8.09 ms after dots onset for monkey H and F, respectively), rising monotonically for the rest of the epoch. The rise in prediction accuracy was steep, reaching 99% (98%) correct for the longest stimuli presentations for monkey H (F), respectively. Moreover, for all 4 epochs considered (targets, dots, delay and post-go) the average accuracy difference between our real-time readout and the equivalent one calculated offline (trained using data from the same session) was within a ± 2% range (Extended Data Fig. 3a-d). Thus, our real-time choice decoder reproduces prediction accuracy as reported in previous off-line analyses of decision-related neural activity in both the oculomotor and somatomotor systems^1, 10^.

Our real-time decoder also reproduced the temporal dynamics and coherence dependence of the DV, as reported in previous off-line studies^1, 10^. The on-line DV: (i) started around 0 at the time of dots onset, (ii) separated by choice after ∼200 ms, and (iii) rose (or fell) faster for easier trials (Fig. 1d, Extended Data Fig. 2b; regression of DV onto coherence significant for both choices, p<10^-5^ uncorrected). Prediction accuracy was higher for correct trials compared to error trials (Extended Data Fig. 4) when holding the stimulus coherence constant, as expected from previous studies^11^.

Finally, our decoding method yielded stable performance across multiple days, justifying combination of data across sessions (Extended Data Fig. 5). This is particularly important when studying rare events such as CoMs, which only happen on a small fraction of the trials and could not be characterized adequately using a single session’s data.

## Real time DV closely predicts choice likelihood across experimental conditions

The previous results are a proof of concept for a highly reliable, real-time readout of decision state in PMd/M1 using spiking data from ∼100-200 units and aggregate and average metrics (Fig. 1c-d, Extended Data Fig. 2a-b). However, we often observed large fluctuations (over 3 natural log units) in the decision variable on individual trials, even within single behavioral epochs (Fig. 1e). If moment-to-moment fluctuations in DV during single trials (as estimated by our decoder) reflect true fluctuations in the decision state of the animal, we expect larger absolute values of DV to be associated with stronger preference for one of the two choices, and hence higher prediction accuracy were a decision to be required at any time during a single trial.

Because we decoded and tracked the DV in real-time, we were able to terminate the visual stimulus in a neurally contingent manner and probe both neural activity and the subject’s behavior with high precision and negligible latency (<34 ms, see Methods). Inspired by sequential sampling behavioral models that assume a bound^12–14^, the first closed-loop test we performed was to impose virtual decision boundaries that, if reached, would result in stimulus termination (Fig. 2a), prompting the subject to immediately report its decision (in trials with no delay period). In this manner we obtained a direct mapping between the nearly instantaneous readout of decision state and the likelihood of a given behavioral choice.

**Figure 2.**
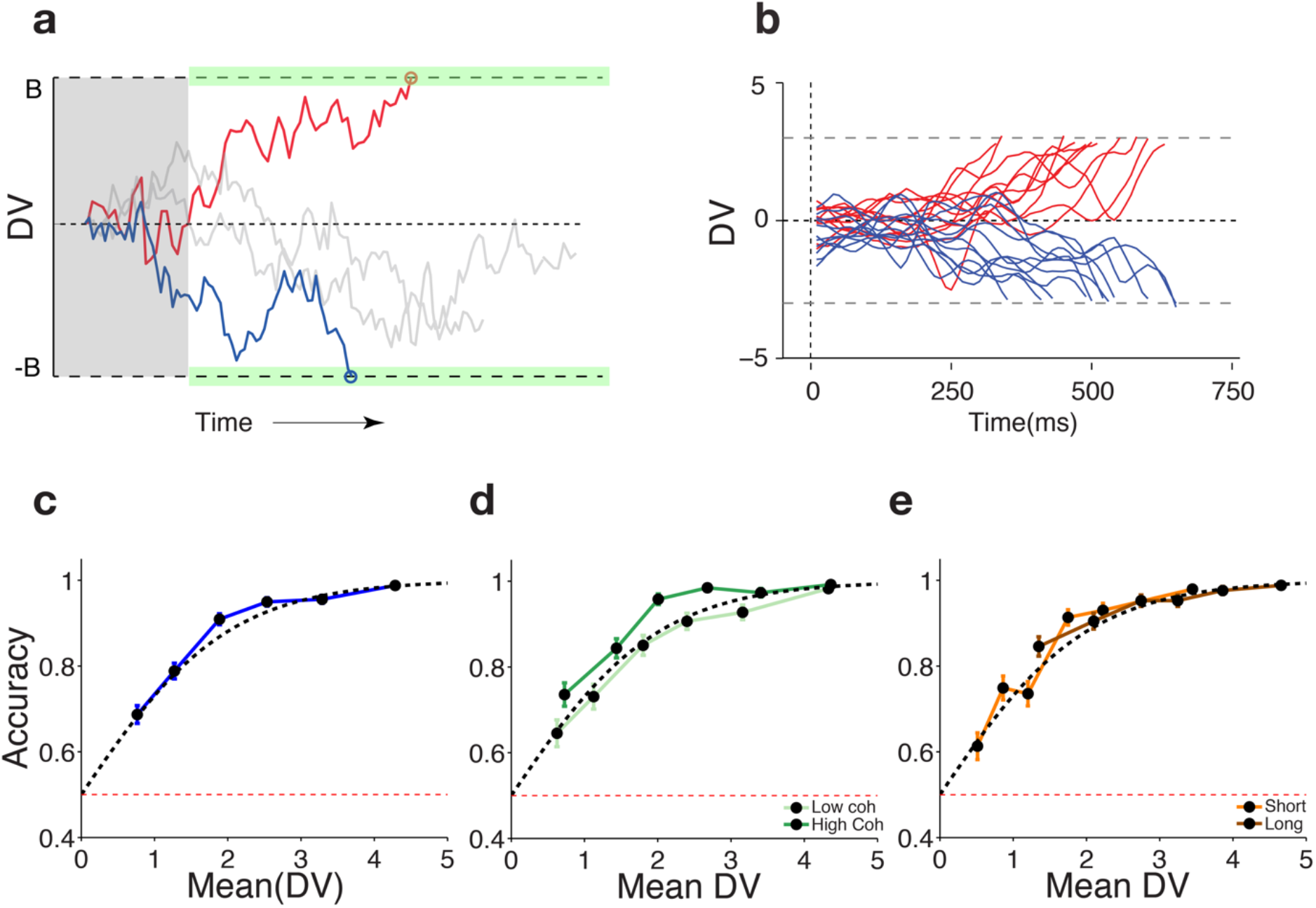
Choice likelihood can be accurately decoded in real-time across experimental conditions using only 50 ms of neural data. **a) Schematic of the first closed loop experiment implemented in real time.** Virtual boundaries for DV magnitude (green shaded regions) were imposed and if reached, triggered the termination of the stimulus presentation. The subject was then immediately asked to report its decision. A 250 ms minimum stimulus duration was imposed (grey shaded region) to prevent random fluctuations in the beginning of the trial from triggering stimulus termination. If the boundary wasn’t reached, the stimulus was presented for a pre-selected random duration (500-1200 ms). Grey traces show cartoons of trials for which the boundary was not reached while red (blue) traces show terminated trials that the decoder predicted would result in a right (left) choice. 5 different boundary values were used on each experiment. **b) Example trials captured during the virtual boundary experiment.** Real-time DV time courses for example trials terminated using boundaries set at +3 and −3 DV units. Traces are colored according to behavioral choice at the end of the trial: right choices in red and left choices in blue. Data from one session from monkey H. **c) Prediction accuracy as a function of DV magnitude.** Choice prediction accuracy for all trials collected during virtual boundary experiment as a function of DV magnitude for monkey H is shown in blue. Trials were split in 6 quantiles sorted by DV magnitude at termination. Prediction accuracy and median DV magnitude were calculated and plotted separately for each quantile (blue line with black symbols). Blue error bars show standard error of the mean for a binomial distribution. Dashed black line shows predicted accuracy from log-odds equation and red dashed line shows chance level. Data from 2973 trials from monkey H. **d) Prediction accuracy as a function of DV and stimulus coherence.** Same data shown in **c)** but having pre-sorted the trials by coherence (see Methods). Dark green trace shows high coherence results and light green, low coherence results. Same conventions as in **c)**. **e) Prediction accuracy as a function of DV and stimulus duration.** Same data shown in **c)** but having pre-sorted the trials by stimulus duration. Brown trace shows results for long trials and orange trace results for short trials (see Methods). Same conventions as in **c)**.

Figure 2b shows 22 example DV traces from trials that led to stimulus termination by reaching a fixed DV boundary of magnitude 3, within a tolerance of ± 0.25 DV units.

To characterize the relationship between the DV at termination and prediction accuracy, we systematically swept the parameter space for the boundary height using values spanning 0.5-5 DV units in 0.5 increments (1DV unit corresponds to an increase of 2.718 in the likelihood ratio of choosing one target over the other). Figure 2c shows that prediction accuracy increases monotonically with the DV magnitude at termination as expected. Moreover, using only 100 ms of data to estimate the DV that triggered termination, the difference between the observed likelihood of a given choice (solid trace) and that predicted by the logistic function (dashed trace) was only, on average, 1.7% (1.9%) for monkey H (F) (Fig. 2c, Extended Data Fig. 2c). For example, neural DV values of ±3 predict decisions upon termination with an accuracy of 98%. Even DV values as low as ±0.5-1 predict decisions with an accuracy of nearly 70%. DV fluctuations below ±0.5 are more susceptible to noise in our estimates of decision state and at most are associated with very weak choice preferences and were thus not tested. Overall, these results show that moment-by-moment fluctuations in PMd/M1 neural population activity captured by our decoding model are indeed reflective of a fluctuating internal decision state of the animal— fluctuations that have been covert and thus uninterpretable until now.

Figure 2c (Extended Data Fig. 2c) combines trials across a wide range of coherences and stimulus durations, aggregated across 17 (15) sessions from monkey H (F). To identify experimental factors that might influence the observed relationship between DV at termination and prediction accuracy, we first resorted the same trials in Figure 3c by stimulus coherence. The results show that there is a small separation between the curves for high and low coherence trials (Fig. 2d) with higher accuracy for high coherence trials. The shift is small but reliable across monkeys (Extended Data Fig. 2d). We hypothesized that this difference resulted from motion energy signals already en route from the retina to PMd/M1 (∼175 ms latency) when the DV reached stimulus termination. More motion energy signals would be arriving from this neural ‘pipeline’ on high coherence trials, leading to a slightly higher DV than we measured at stimulus termination.

**Figure 3.**
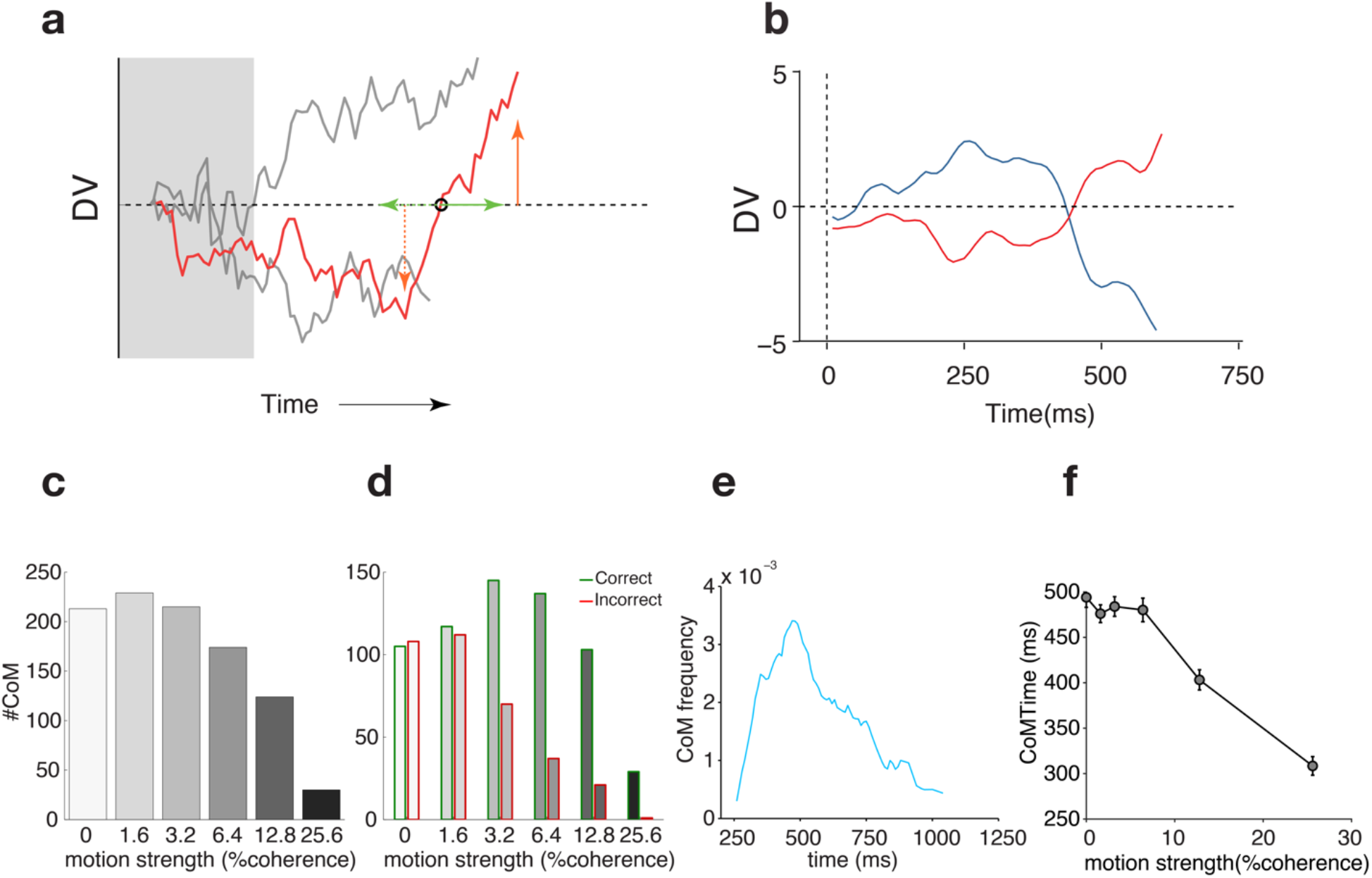
Putative changes of mind can be detected and validated in real time. **a) Schematic of the second closed loop experiment implemented in real time.** The value and history of the DV trace were tracked on each trial. If a 0 crossing (sign change in the DV) was detected, the conditions required for termination were checked and termination was carried out if the conditions were met (see Methods). In this example the conditions for temporal stability of DV sign are depicted by the green horizontal arrows while the conditions for minimum DV deflection before and after CoM are depicted by the orange arrows. Upon termination, the subject was immediately asked to report its decision. A 250 ms minimum stimulus duration was imposed (grey shaded region) such that random fluctuations in the beginning of the trial did not trigger stimulus termination. If the conditions were not met or if a 0 crossing was never detected, the stimulus would be presented for a pre-selected random duration (500-1200 ms). Grey traces show cartoons of trials for which the 0 crossings would not meet the criteria while the red the trace shows a terminated trial that was predicted to lead to a rightward choice. One set of criteria for CoM validity was used in each session (Extended Data. Table 4). **b) Example trials captured during the CoM experiment.** Real-time DV time courses for 2 example trials with a putative CoM terminated after conditions were met (minimum DV pre and post CoM: 2 and minimum period of sign stability pre and post CoM: 150 ms). Traces are colored according to behavioral choice at the end of the trial: right choices in red and left choices in blue. Two trials from one session from monkey H. **c) CoM frequency as a function of coherence**. Total number of CoMs detected for each coherence for monkey H. **d) CoM frequency as a function of coherence and direction**. Total number of CoMs detected for each coherence and direction for monkey H. Red bars correspond to erroneous CoMs and green bars to corrective CoMs. **e) CoM frequency as a function of time in the trial**. Frequency of CoMs detected as a function of time during stimulus presentation for monkey H. Because only CoMs that would have resolved by 250 msec after stimulus onset were considered, there is an edge effect with CoM frequency briefly increasing between ∼250-450 msec after which it declines. **f) CoM time as a function of coherence.** Average CoM time (defined as the zero crossing for each CoM trial) is plotted as a function of stimulus coherence. Error bars show s.e.m across trials for each condition. CoM time was negatively correlated with stimulus coherence (p =1.8 x 10^-17^)

To assess this possibility, we measured the derivative of the DV around termination and performed the following two analyses. First we checked whether DV derivative explained a significant fraction of choice variance beyond DV value alone (see Methods). For both monkeys the effect of DV derivative (defined as the DV slope in the last 50 ms of stimulus presentation) was significant (p=0.02, p = 4.5×10^-11^ for monkey H and F, respectively) and the effect was congruent with our hypothesis: stronger positive derivatives predicted higher likelihood of rightward choices and stronger negative derivatives predicted higher likelihood of leftward choices (Extended Data Table 1, “DV diff”). Second, we tested whether high coherence trials were associated with higher DV derivatives at termination by performing linear regression of DV derivatives as a function of signed coherence. For both monkeys signed coherence was strongly predictive of DV slopes: p = 2.17 x10^-171^ and R^2^ = 0.23 for monkey H and p = 1.57 x10^-10^^5^ and R^2^ = 0.16 for monkey F. These results confirm that DV derivative is predictive of choice beyond DV alone and show that higher coherence trials are associated with higher DV derivatives. The data are consistent with our hypothesis above that the DV continues to evolve under the influence of ‘pipeline’ sensory information for a short interval following stimulus termination, resulting in somewhat better prediction accuracy than expected from the DV at termination, especially at high coherences.

Sorting trials by duration (Fig. 2e, Extended Data Fig. 2e), reveals a different effect: the centers of the quantiles are strongly shifted to the right (higher DV magnitudes) for longer stimuli compared to shorter stimuli. This effect is expected from multiple sequential sampling models^8,15–17^. In drift diffusion models, for example, diffusion to high decision bounds requires more time than for low bounds^18^. However, we tested whether stimulus duration *per se* was a significant predictor of choice independently of DV value by including two additional regressors in our logistic model of choice: stimulus duration (representing choice bias as a function of time) and an interaction term between stimulus duration and direction (representing increased sensitivity to stimulus coherence as function of time). Neither regressor was significant for either monkey (p>0.05, Extended Data Table 1), implying that the likelihood of making one or the other choice depended on DV value independently of the time required to reach that value.

Together, these results show that fluctuations in DV magnitude at a 100 ms time scale have a predictable correlate in choice likelihood that is lawfully influenced by stimulus coherence and robust across time. We emphasize that our decoded DV is model-based and thus a proxy for the actual decision state in the brain. We are sampling from a relatively small number of neurons, and the underlying mechanism is unlikely to be strictly linear (in contrast to the logistic model). In addition, we do not know with certainty when the deliberation process ends within the brain, which could occur before or after our stimulus termination on individual trials. Despite these caveats, our ability to predict choice likelihood using a DV boundary criterion at stimulus termination within a very small margin of error (<2% on average) confirms that DV is a reliable proxy for decision state.

## Neurally detected CoMs can be validated and match the statistical regularities expected from previous studies

The mapping between DV and choice likelihood obtained in the first experiment (Fig. 2c), enabled us to perform a new closed-loop experiment aimed at capturing particularly robust DV fluctuations in which the sign of the DV (and thus the neurally inferred decision state of the animal) changed in the middle of a trial, suggestive of a ‘change of mind’ at the behavioral level (CoM, Fig. 3a-b). When the neural criteria for a CoM were met in real-time (see Methods, examples in Figure 4a, orange and green arrows), the stimulus was terminated instructing the monkey to make a decision as described above. Our aim was to detect neurally-based candidate CoMs, assess the influence of the decision states before and after the CoM on the final choice, and determine whether statistical properties of the neurally derived CoMs match the properties expected of CoMs from prior psychophysical and neurophysiological studies.

**Figure 4.**
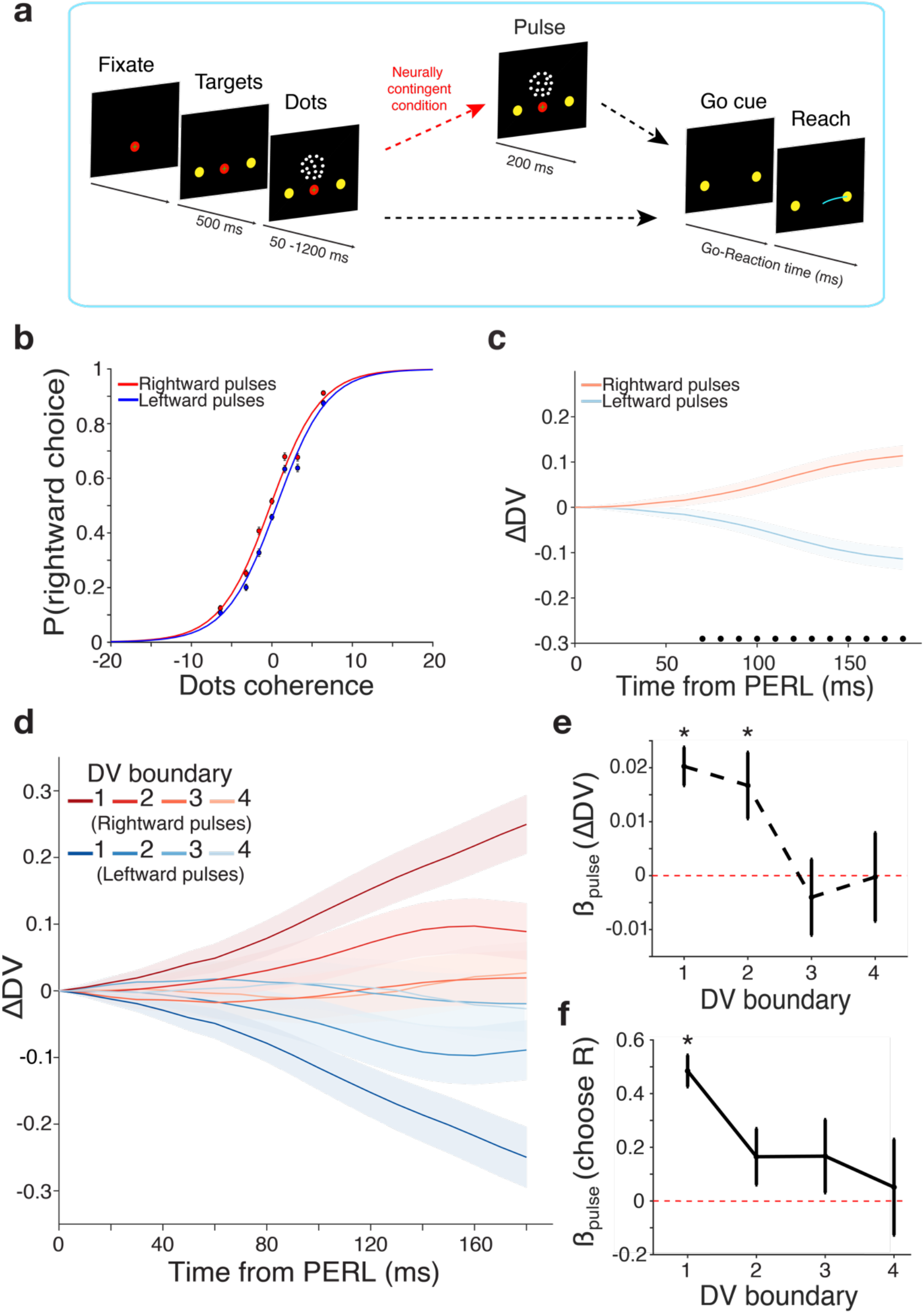
Neurally triggered pulses of motion evidence nonlinearly bias both choice behavior and DV. **a) Motion pulse task.** As in the motion discrimination task, trials began with the onset of a fixation point (FP) on the touchscreen. Once both eye and hand fixation were acquired, two targets appeared. The motion stimulus was shown after a short delay (500 ms) and a maximum stimulus duration was randomly assigned from 500-1200 ms. Virtual boundaries for DV magnitude were imposed (randomly assigned to integer values from 1-4) and if reached, triggered a 200-ms pulse of additive dots coherence, randomly assigned to be rightward or leftward on each trial (± 2% coherence for monkey H), followed immediately by termination of the dots stimulus. A 50 ms minimum stimulus duration was imposed to ensure a minimum total stimulus duration of 250 ms. If the DV boundary wasn’t reached, the dots stimulus was presented for a pre-selected random duration (500-1200 ms). Dots offset was followed by the go cue. Decision states were continuously decoded using the dots period decoder during all epochs of the trial (blue box, see Methods). **b) Psychometric functions for pulse trials.** Curves were fit using logistic regression on choice with signed stimulus coherence and pulse direction as predictors, plus an intercept term. Data points show mean proportion of rightward choices for each stimulus coherence, ± s.e.m. The pulse effect is equivalent to changing the overall stimulus coherence by 0.384% (standard error 0.0514%, p = 8.38E-14). Data from 9614 rightward and 9523 leftward pulse trials from monkey H. **c) Average change in post-pulse DV from estimated Pulse Evidence Representation Latency (PERL), mean subtracted.** ΔDV is the difference in the DV at each time point from the DV at the PERL (170 ms for monkey H; see Methods). The mean ΔDV across pulse directions in each time bin has been subtracted for visualization. Shaded error bars correspond to mean ± s.e.m. Black dots indicate time bins in which ΔDV is significantly different for trials with pulses in opposite directions (false discovery rate 0.05). Data from same trials as **b)**. **d) Average change in post-pulse DV for each DV boundary, mean subtracted.** Conventions as in **b)** but separated by the DV boundary triggering motion pulses in each direction. Darker colors correspond to smaller DV boundary magnitudes. Data from monkey H, minimum 1507 trials per condition shown. **e) Pulse coefficients from linear regression on ΔDV slope for each DV boundary.** ΔDV slope is the single-trial slope of the ΔDV from PERL to either the animal’s median go-reaction time or 150 ms prior to movement onset, whichever came first (as shown in **b)**). Multiple linear regression was performed separately on ΔDV slope for trials at each DV boundary with the following predictors: signed dots coherence, pulse onset time, pulse direction, pulse onset time * pulse direction, plus an intercept term. Data points and error bars represent the coefficient for pulse direction for trials at each DV boundary, ± s.e.m.; asterisks denote significantly nonzero coefficients at 95% confidence. Data from same trials as **d)**. **f) Pulse coefficients from logistic regression on choice for each DV boundary.** Logistic regression was performed separately on the probability of a rightward choice for trials at each DV boundary with the following predictors: signed dots coherence, pulse onset time, pulse direction, pulse onset time * pulse direction, plus an intercept term. Data points and error bars represent the coefficient for pulse direction for trials at each DV boundary, ± s.e.m.; asterisks denote significantly nonzero coefficients at 95% confidence. Data from same trials as **d)**.

We conceptually divide a CoM trial into two segments—the initial preference prior to the DV sign change, and the final (opposite) preference that leads to the observed choice. The observed choices allow corroboration of the neural estimate of the final decision state in the second segment (Extended Data Fig. 6). For monkey F, the relationship between choice prediction accuracy and DV at stimulus termination for CoM trials was very similar to that of non-CoM trials (compare Extended Data Fig. 2c and Extended Data Fig. 6b, mean error between predicted and observed choice likelihood: 1.9% for non-CoM trials vs 3.8% for CoM trials). This relationship was lawful and monotonic for monkey H as well although lower than expected (Extended Data Fig. 6a, compared to Fig. 2c, mean error between predicted and observed choice likelihood: 1.7% for non-CoM trials vs 9.3% for CoM trials), suggesting that in addition to the measured DV at stimulus termination, monkey H’s decisions were also influenced by some aspect of the DV trajectory history specifically related to the CoM. We formally tested this hypothesis by regressing choice as a function of 3 additional parameters (in addition to DV at termination) that were enforced and monitored in this experiment (see Methods): maximum DV deflection before sign change and duration of sign stability before and after DV sign change. For monkey F, no additional factor was choice predictive, whereas for monkey H both the duration of sign stability before and after the CoM were also choice predictive (Extended Data Table 2) as suspected from Extended Data Fig. 6.

We combined all 985 (1727) CoM’s detected in monkey H (F) to assess whether our neurally detected CoMs conformed to three statistical regularities of CoMs established in previous psychophysical^7^ and electrophysiological^1^ studies.

The first observation is that CoMs are more frequent for low and intermediate coherence trials as opposed to high coherence trials, as high coherences are more likely to lead to straightforward integration of evidence toward the correct choice. We found the same to be true in our real-time detection data (Fig. 3c, Extended Data Fig. 2f; linear regression p<0.001).

The second observation is that CoMs are more likely to be corrective than erroneous. This prediction results from the corrective role of additional visual evidence on the initial preference of the subjects. This trend was also verified in the CoMs we detected with CoMs for non-zero coherences and for both monkeys being more likely corrective than erroneous (Fig. 3d, Extended Data Fig. 2g; Wilcoxon rank sum test p<0.001, median corrective and erroneous CoM counts: 530 and 242 for monkey H and 1046 and 443 for monkey F, respectively).

Finally, the third observation made in these previous studies was that CoMs were more frequent early in the trial than later in the trial, consistent with drift diffusion models in which the DV is more likely to have hit an absorbing decision bound as the trial progresses. We observed this effect in our real-time, neurally detected CoMs as well (Fig. 3e, Extended Data Fig. 2h).

We also discovered a new regularity associated with CoMs: the average time of zero crossing was negatively correlated with stimulus coherence (Fig. 3f, Extended Data Fig. 2i). This observation likely results from the stronger corrective effect of higher coherence stimuli (Fig. 3d, Extended Data Fig. 2g).

Together, these results show that robust fluctuations in DV that imply a change in choice preference (zero crossing) can be captured in real time and validated as changes of mind.

## Pulses of additional visual motion evidence have smaller neural and behavioral effects when presented at larger DV values

In a final set of closed-loop experiments whether the neural and behavioral responses to brief pulses of additional motion information varied with the state of the DV before the pulse. Inspired by decision-making models involving buildup of neural activity to a bound^15, 16, 19, 20^, we expected termination of the deliberation process and commitment to a choice to be more likely at high DV values^1, 7, 8, 16, 21^. We therefore hypothesized that additional pulses of sensory evidence would result in less change in DV and behavior when pulses were triggered by high DV values.

To characterize the relationship between DV and responses to a stimulus pulse, we again imposed virtual DV boundaries (as in Fig. 3a-b) that, if reached, triggered a 200-ms pulse of additive dots coherence (randomly assigned to be rightward or leftward on each trial) followed by stimulus termination (Fig. 4a). We swept a subset of the previously used DV values for the boundary (spanning 1-4 DV units, in 1.0 increments). Pulse strength was calibrated to yield very small but significant effects on behavior, in an effort to avoid making the pulses so salient as to change the animals’ integration strategy on pulse trials (Δcoherence = 2% for monkey H, 4.5% for monkey F). Pulse information had no bearing on the reward^8, 17^. Motion pulses slightly but significantly biased the monkeys’ choices in the direction of the pulse (p = 8.38E-14 for monkey H, Fig. 4b; p = 1.95E-4 for monkey F, Extended Data Fig. 2j).

We reasoned that, to detect the presumably small effects of these small motion pulses on the DV, we would need to account for a processing delay for changing stimulus information to influence our recorded neural populations. Thus, to quantify the effect of the pulse on the evolving DV, we first measured the minimum latency for visual stimulus information to influence the DV: we calculated the time after stimulus onset at which the DV traces diverged for rightward vs. leftward choices in an independent set of open loop trials at the strongest motion coherence. We refer to this time point as the evidence representation latency (ERL). For each trial, we measured the change in DV (ΔDV) for each time bin, beginning at the time of pulse onset plus the ERL (or PERL—see Methods). We found that, on average, motion pulses slightly but significantly biased ΔDV in the direction of the pulse (Fig. 4c, Extended Data Fig. 2k).

In the case of simple, unbounded linear integration, we expect the magnitude of DV change in response to a fixed motion pulse to remain constant regardless of the triggering DV at pulse onset. In contrast, Fig. 4d (Extended Data Fig. 2l) shows that motion pulses led to larger DV changes when triggered by low as compared to high DV values.

Previous studies have shown that behavioral and LIP neural responses to similar motion pulses tend to be smaller when pulses are delivered later in the stimulus^8, 17^. Large DV values tend to occur later in the trial, and this was hypothesized to be the underlying reason for the diminishing pulse effects (assuming some sort of bound on integration of evidence at larger DVs); but these studies lacked concurrent neural population recording and decoding and thus did not have access to the momentary decision state. Thus, the time of pulse onset is a possible confound for the decreasing pulse effects at high DV bound values as depicted in Fig 4d. To control for this possibility, we first used the slope of the ΔDV vs. time relationship measured on individual trials (ΔDV slope) to summarize the effect of the stimulus pulse on DV on single trials. We then performed a multiple regression analysis of ΔDV slope that included both the triggering DV value and the time of the motion pulse as regressors (and other variables as well - see Methods). The regression data confirm that the effect of stimulus pulses on DV is only significant when triggered by lower DV values, and that this effect is not explained by pulse timing (Fig. 4e, Extended Data Fig. 2m, Supplementary Information Table 1). Similarly, the effect of motion pulses on psychophysical behavior is weaker when triggered by high DV values, and these effects also are not explained by pulse timing (Fig. 4f, Extended Data Fig. 2n, Supplementary Information Table 2).

Our finding that larger DVs (and corresponding behavioral readouts) are more resistant to pulses is consistent with several models of decision formation, including linear integration to a decision bound (such as a simple stopping criterion^12^) or a more complex nonlinear integration process^17, 22, 23^. Inspired by these results, we returned to the data from the first two experiments in an attempt to explore the nature of the apparent bounding mechanism by analyzing the time-variance of the DV over the course of individual trials. In the case of an absorbing decision bound or attractor network, we would expect DV variability (measured as the DV time derivative) to decrease after reaching the bound. We indeed found that, on average, DV variability decreases over time within single trials (Fig. 5a, Extended Data Fig. 7a). This effect holds across all stimulus strengths, although variability peaks earlier and falls faster on the easiest trials (Fig. 5b, Extended Data Fig. 7b).

**Figure 5.**
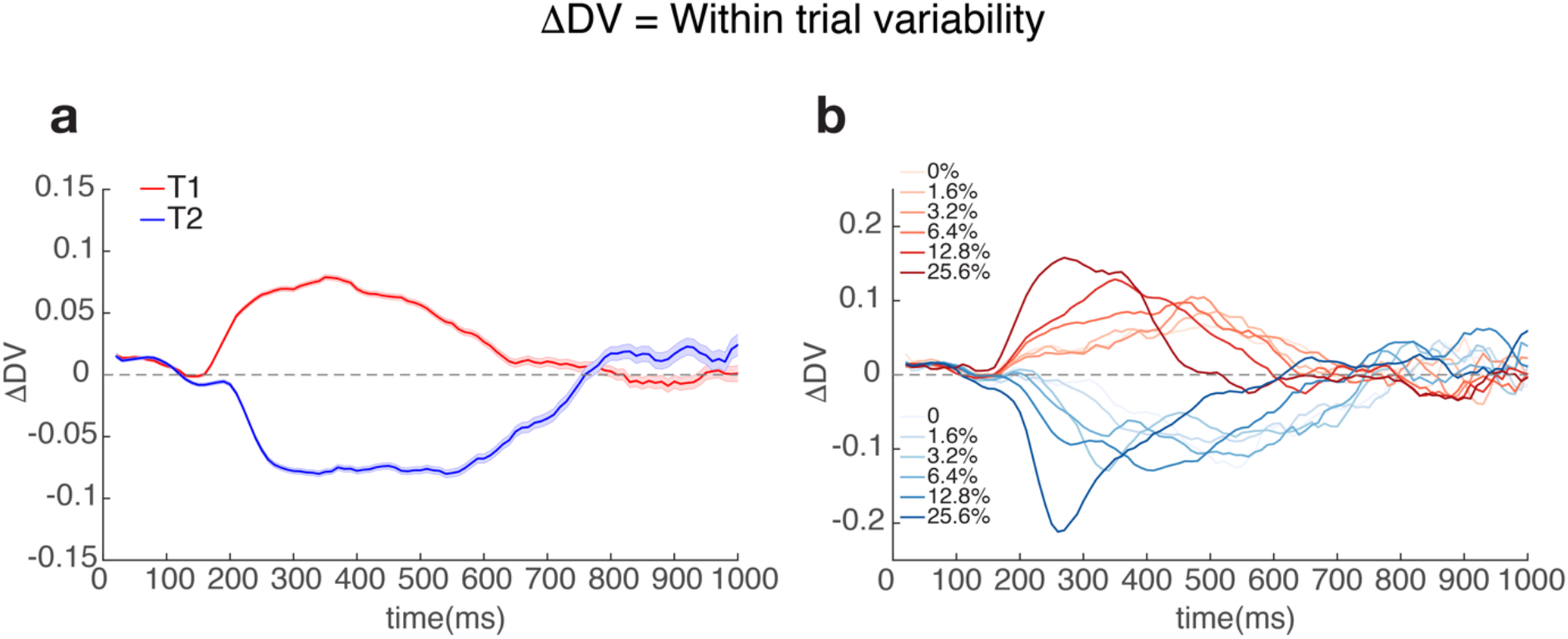
Within trial DV variability decreases over time for long duration stimuli. **a) Average DV derivative as a function of time and choice - monkey H.** ΔDV was calculated for each trial as the difference between consecutive DV estimates spaced out by 10 ms. Traces show average ΔDV +/-s.e.m for right choices (red trace) and left choices (blue trace) during stimulus presentation. ΔDV initially starts increasing around the expected stimulus latency (170 ms) but progressively decreases for long (>600 ms) stimulus presentations. **b) Average DV derivative as a function of time and signed coherence - monkey H.** Same data as in **a)** but with DV derivative averaged separately for each choice and motion coherence level (correct trials only). Right choices are plotted in red and left choices in blue as in **a)**. Darker traces correspond to stronger coherences.

## Discussion

While previous single-electrode recordings have strongly advanced our understanding of the neural correlates of perceptual decision-making, interesting dynamics in choice signals were lost to necessity of averaging data across trials. With a few notable exceptions^3, 24^, deploying the statistical power of simultaneous multi-electrode recordings to track single-trial population dynamics during choice behavior is a recent advance^1, 2, 6^. Even more recent work has leveraged the power of brain-computer interfaces (BCIs) to study neural correlates of prediction, learning, and multisensory integration (as reviewed in Golub et al. 2016^25^). In this study, for the first time, we probed moment-to-moment fluctuations in decision states using BCI-inspired closed loop experiments that enabled neurally contingent stimulus control and made behavioral validation of these fluctuations feasible (see Methods). We show that large fluctuations (up to several log units) in a decoded decision variable in premotor and primary motor cortices are nearly instantaneously (<100 ms) predictive of choice. We captured neural correlates of changes of mind in the form of robust changes in DV sign. The statistical regularities of these rare events match previous psychophysical CoM findings. Finally, we showed that larger DV values are resistant to additional pulses of sensory evidence, supporting the hypothesis that large DVs are associated with higher commitment to an upcoming choice.

Importantly, the impressive choice prediction accuracy achieved in this study using a linear decoder does not imply that the brain’s decision formation process is also linear. In principle, such a decoder could predict binary choices quite well even if the true neural process underlying decision formation were nonlinear, depending on the form of the nonlinearity (see, e.g., Sussillo et al. 2016^26^ for an example of a linear neural to kinematic decoder which only slightly underperforms a more powerful nonlinear recurrent neural network). However, our linear DV is tightly linked to choice behavior (e.g. Fig. 2c), showing that variations in DV magnitude meaningfully track the ongoing process of decision formation despite the possible presence of nonlinearities in the underlying neural mechanism.

Previous studies have described fluctuations in offline decoded decisions associated with changes of mind^1–3, 6^. Here we confirm and extend those observations with neurally contingent interrogation of candidate CoM events, but we also find large, behaviorally relevant fluctuations even when the DV remains on one side of the discriminant hyperplane in non-CoM trials (e.g. Figs. 1e and 2b). We wondered whether these DV fluctuations were related to stochastic variations in motion strength of the stimulus on single trials. While across coherence levels the average motion energy explains a large portion of DV variance (Extended Data Fig. 8a-b), our data shows that within coherence stochastic fluctuations in the stimulus are not the dominant cause of DV fluctuations (Extended Data Fig. 8c-d). Further experiments will be needed to address the source(s) of these fluctuations and their relationship with fluctuations in other brain areas^27^ as well as other cognitive processes including motor preparation and execution^28, 29^, attention, motivation, and confidence.

In addition to validating the behavioral relevance of neurally detected DV fluctuations, our ability to impose real-time task changes contingent upon them allowed us to show that neural and behavioral responses to pulses of additional sensory evidence diminish when pulses are presented at larger momentary DV values. These results, combined with the reduction in DV variability observed over the course of single trials, suggest the presence of an absorbing decision bound in these motor cortical neural populations, consistent with attractor dynamics in which the neural population converges on a stable state as a decision is formed^22, 23^.

The conceptual and technical innovation that enabled these findings is our ability to accurately decode decision states in real time, which could bring the concept of cognitive prostheses^30–33^ much closer to reality by providing another means of decoding subjects’ goals for use as a flexible prosthetic control signal. More broadly, the real-time closed loop approach demonstrated here may be applicable not only to decision-making processes, but also to other cognitive phenomena such as working memory and attention.

## Extended Data

**Extended Data Table 1.** Coefficients obtained from logistic regression on choice – virtual boundary experiment (monkeys H and F)

| Logistic Regression on Choice |  |  |  |  |  |  |
| --- | --- | --- | --- | --- | --- | --- |
|  | Monkey H |  |  | Monkey F |  |  |
| Predictor | Beta Value | 95% CI | p-value | Beta Value | 95% CI | p-value |
| Bias | -0.1613 | [-0.2987 , -0.02381] | 0.02147 | -0.1197 | [-0.2413 , 0.001806] | 0.0535 |
| Coherence | 2.747 | [ 2.268 , 3.227] | 2.757e-29 | 1.885 | [ 1.55 , 2.22] | 2.912e-28 |
| DV Termination | 2.073 | [ 1.779 , 2.367] | 1.887e-43 | 1.708 | [ 1.52 , 1.895] | 1.626e-71 |
| DV Diff | 0.2989 | [0.0488 , 0.5491] | 0.01916 | 0.577 | [0.4062 , 0.7478] | 3.586e-11 |
| Stimulus duration | 0.1199 | [-0.01489 , 0.2548] | 0.08125 | 0.07474 | [-0.04713 , 0.1966] | 0.2293 |
| Stimulus duration*<br>Stimulus direction | -0.02803 | [-0.2053 , 0.1492] | 0.7566 | 0.0466 | [-0.1132 , 0.2064] | 0.5677 |

**Extended Data Table 2.** Coefficients obtained from logistic regression on choice - change of mind experiment (monkeys H and F)

| Logistic Regression on Choice after CoM |  |  |  |  |  |  |
| --- | --- | --- | --- | --- | --- | --- |
|  | Monkey H |  |  | Monkey F |  |  |
| Predictor | Beta Value | 95% CI | p-value | Beta Value | 95% CI | p-value |
| Bias | -0.2876 | [-0.4658 , -0.1093] | 0.001568 | -0.192 | [-0.3773 , -0.00662] | 0.04235 |
| Coherence | 1.304 | [ 1.005 , 1.602] | 1.189e-17 | 1.284 | [ 0.96 , 1.608] | 8.028e-15 |
| DV Termination | 1.67 | [ 1.158 , 2.182] | 1.623e-10 | 2.46 | [ 1.872 , 3.049] | 2.51e-16 |
| DV max opposite | 0.06967 | [-0.4461 , 0.5854] | 0.7912 | -0.2387 | [-0.8959 , 0.4185] | 0.4766 |
| Time after CoM *<br>sign(DV<br>Termination) | 0.7065 | [0.2317 , 1.181] | 0.003544 | 0.02265 | [-0.3835 , 0.4288] | 0.913 |
| Time before CoM<br>* sign(DV max<br>opposite) | 0.7471 | [0.3083 , 1.186] | 0.0008467 | -0.1427 | [-0.6394 , 0.3539] | 0.5732 |

**Extended Data Figure 1.**
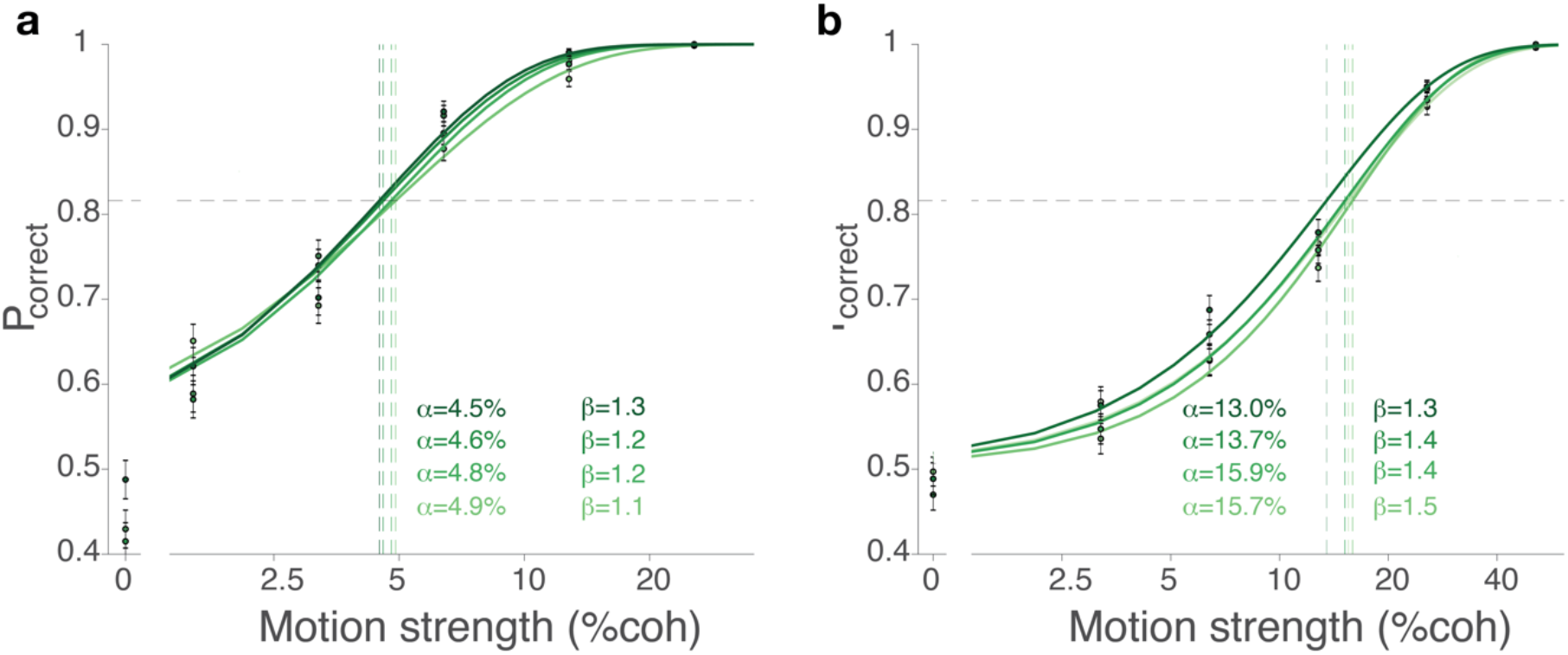
Behavioral performance - Variable duration task. **a) Psychophysical performance for monkey H in the variable duration task.** Percentage correct is plotted as a function of net motion coherence (calculated for both directions). Trials were sorted for stimulus duration in 4 quartiles from long (dark green curve) to short (light green curve). Data from each quartile were fit separately by a Weibull curve. Inset shows fit parameters for each quartile. Data from 12516 open loop trials. Stimulus duration quartiles: Q1: [0.500, 0.574] s Q2: [0.574, 0.680] s Q3[0.680, 0.827] s Q4: [0.827, 1.200] s. **b) Psychophysical performance for monkey F in the variable duration task.** Same as **a)** for monkey F. Data from 12365 open loop trials. Stimulus duration quartiles: Q1: [0.500, 0.574] s Q2: [0.574, 0.667] s Q3[0.667, 0.813] s Q4: [0. 813, 1.200] s.

**Extended Data Figure 2.**
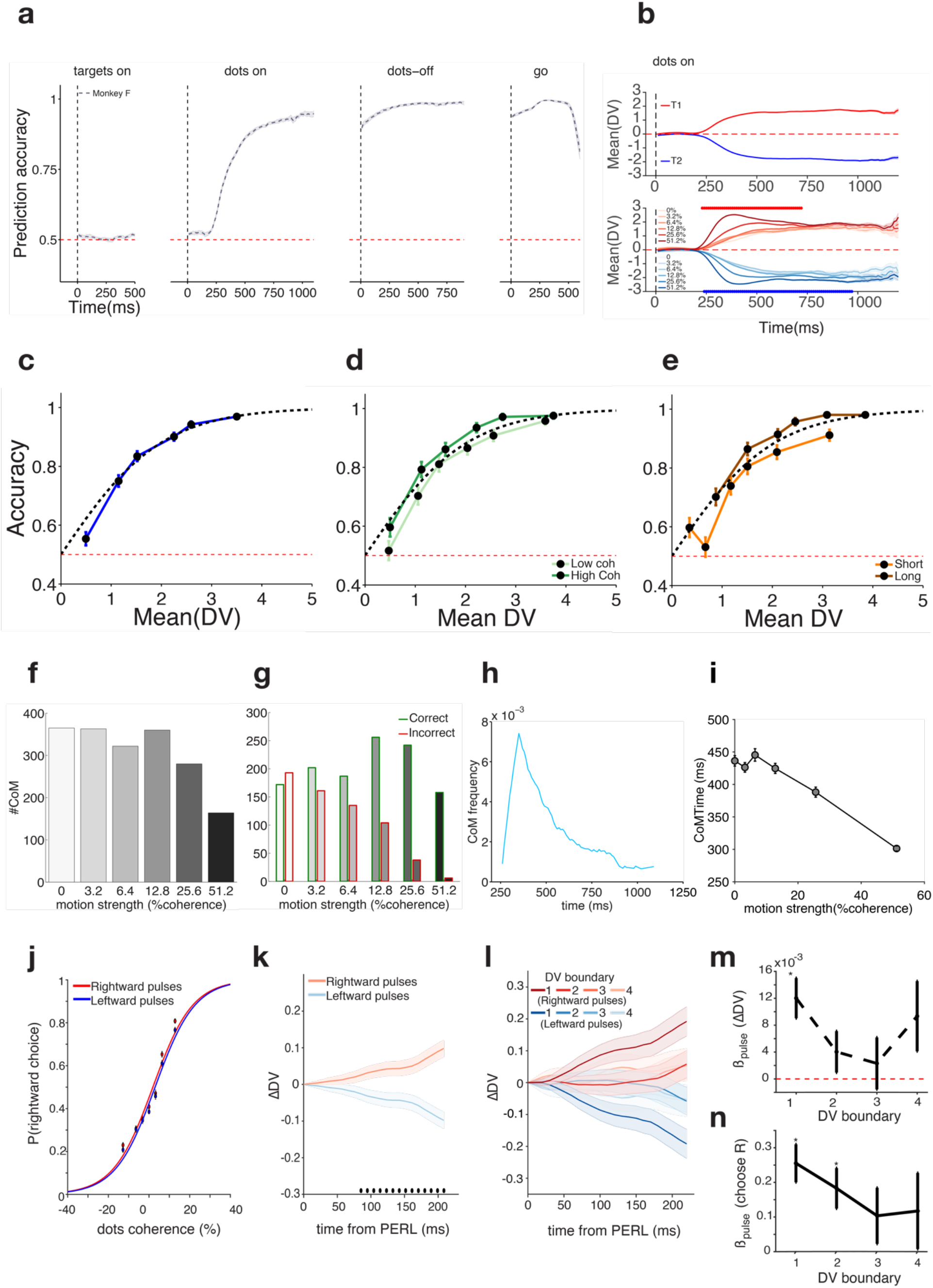
Results for decoding and perturbation of DV in real time - monkey F. **a) Choice prediction accuracy obtained from real-time readout.** Same as Figure 1c for monkey F. Accuracy was calculated for each session and averaged across sessions using a total of 15826 trials. **b) Average Decision Variable traces during dots period.** Same as Figure 1d for monkey F. For monkey F coherence is a significant regressor of DV for at least one of the choices for the period between [230, 970] ms aligned to dots onset. **c) Prediction accuracy as a function of DV magnitude.** Same as Figure 2c for monkey F. Data from 2518 trials from monkey F. **d) Prediction accuracy as a function of DV and stimulus coherence.** Same data shown in **c** but having pre-sorted the trials by coherence (see Methods). Dark green trace shows high coherence results and light green, low coherence results. Same conventions as in **c**. **e) Prediction accuracy as a function of DV and stimulus duration.** Same data shown in **c)** but having pre-sorted the trials by stimulus duration. Brown trace shows results for long trials and orange trace results for short trials (see Methods). Same conventions as in **c)**. **f) CoM frequency as a function of coherence**. Same as Figure 3c for monkey F. **g) CoM frequency as a function of coherence and direction**. Same as Figure 3d for monkey F. **h) CoM frequency as a function of time in the trial**. Same as Figure 3e for monkey F. **i) CoM time as a function of coherence.** Same as Figure 3f for monkey F. CoM time was negatively correlated with stimulus coherence (p =3.0x 10^-30^) **j) Psychometric functions for pulse trials.** Same as Figure 4b for monkey F. The pulse effect is equivalent to changing the overall stimulus coherence by 0.545% (standard error 0.146%, p = 1.95E-4). Data from 10370 rightward and 9967 leftward pulse trials. **k) Average change in post-pulse DV from estimated Pulse Evidence Representation Latency (PERL), mean subtracted.** Same as Figure 4c for monkey F. PERL = 180 ms. Data from same trials as **j)**. **l) Average change in post-pulse DV for each DV boundary, mean subtracted.** Same as Figure 4d for monkey F. Minimum 1731 trials per condition shown. **m) Pulse coefficients from linear regression on ΔDV slope for each DV boundary.** Same as Figure 4e for monkey F. **n) Pulse coefficients from logistic regression on choice for each DV boundary.** Same as Figure 4f for monkey F.

**Extended Data Figure 3.**
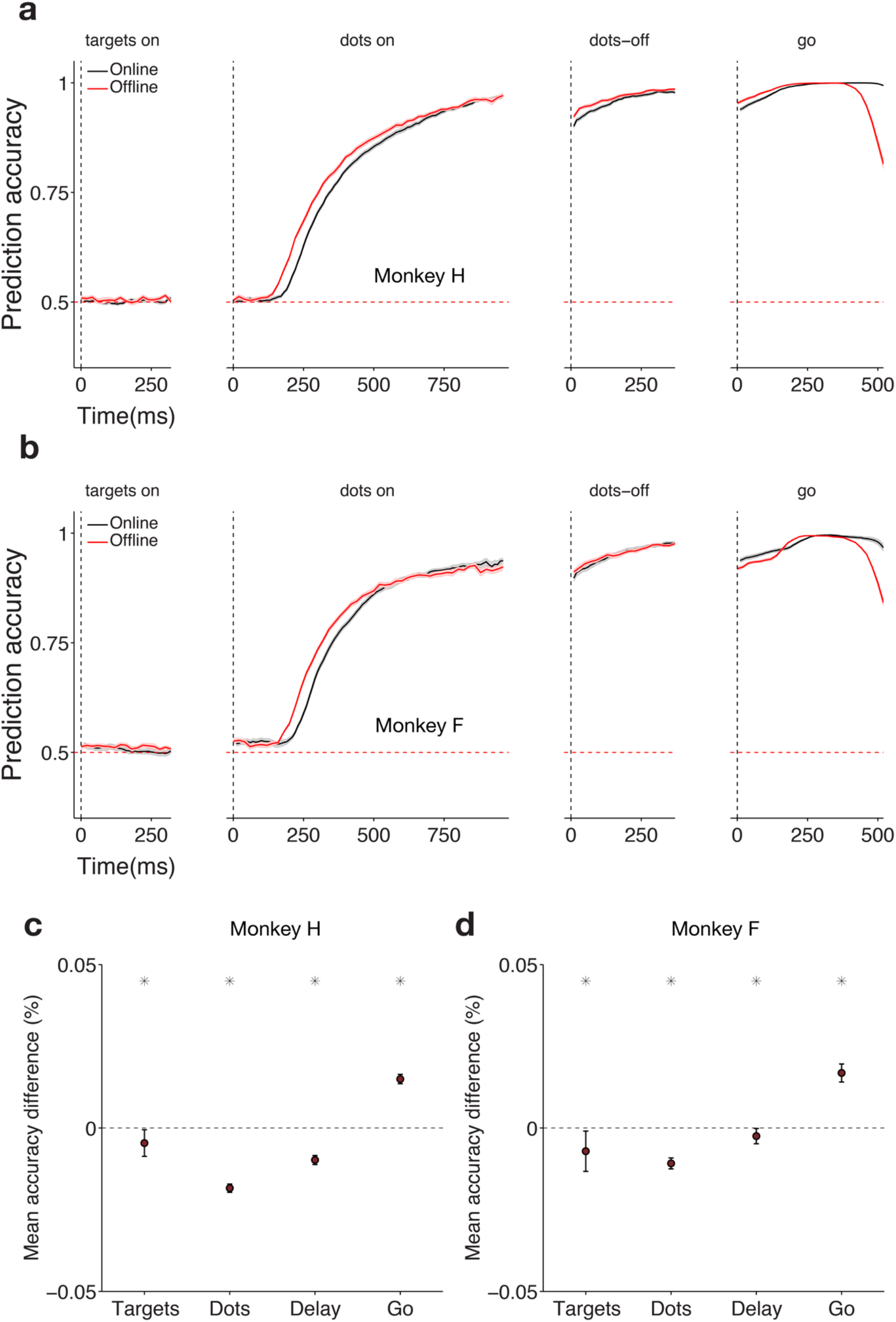
Prediction accuracy online, offline and as a function of stimulus coherence. **a) Online and Offline classifiers result in similar performance for targets, dots delay and post-go epochs for monkey H.** Average prediction accuracy (see Methods) over time ± SEM (across sessions) for monkey H. Online / offline classifier results are plotted in black / red. Data in black is re-plotted from Figure 2a. Prediction accuracy is very similar online and offline across the trial (see **c)**). **b) Online and Offline classifiers result in similar performance for targets, dots delay and post-go epochs for monkey F.** Same as **a** but for monkey F. Same conventions apply. **c) Summary of performance difference between online and offline classifiers within each epoch for monkey H.** Average performance difference between online and offline classifiers (accuracy difference in percentage correct) for each of the epochs plotted in a). Positive number numbers correspond to better online classifier performance and negative numbers to better offline classifier performance. Black asterisks correspond to windows for which the differences were significantly larger than zero (Wilcoxon signed-rank test, P<0.001). **d) Summary of performance difference between online and offline classifiers within each epoch for monkey F.** Same as **c)** for monkey F. For both monkeys **c)** and **d)** the difference of choice prediction accuracies between the online and the offline classifiers was small and negative for the target, dots and delay epochs (between −0.2% and −1.9%). In contrast, for the post-go period, the difference in prediction accuracies was slightly positive (1.5% and 1.7% for monkey H and F respectively).

**Extended Data Figure 4.**
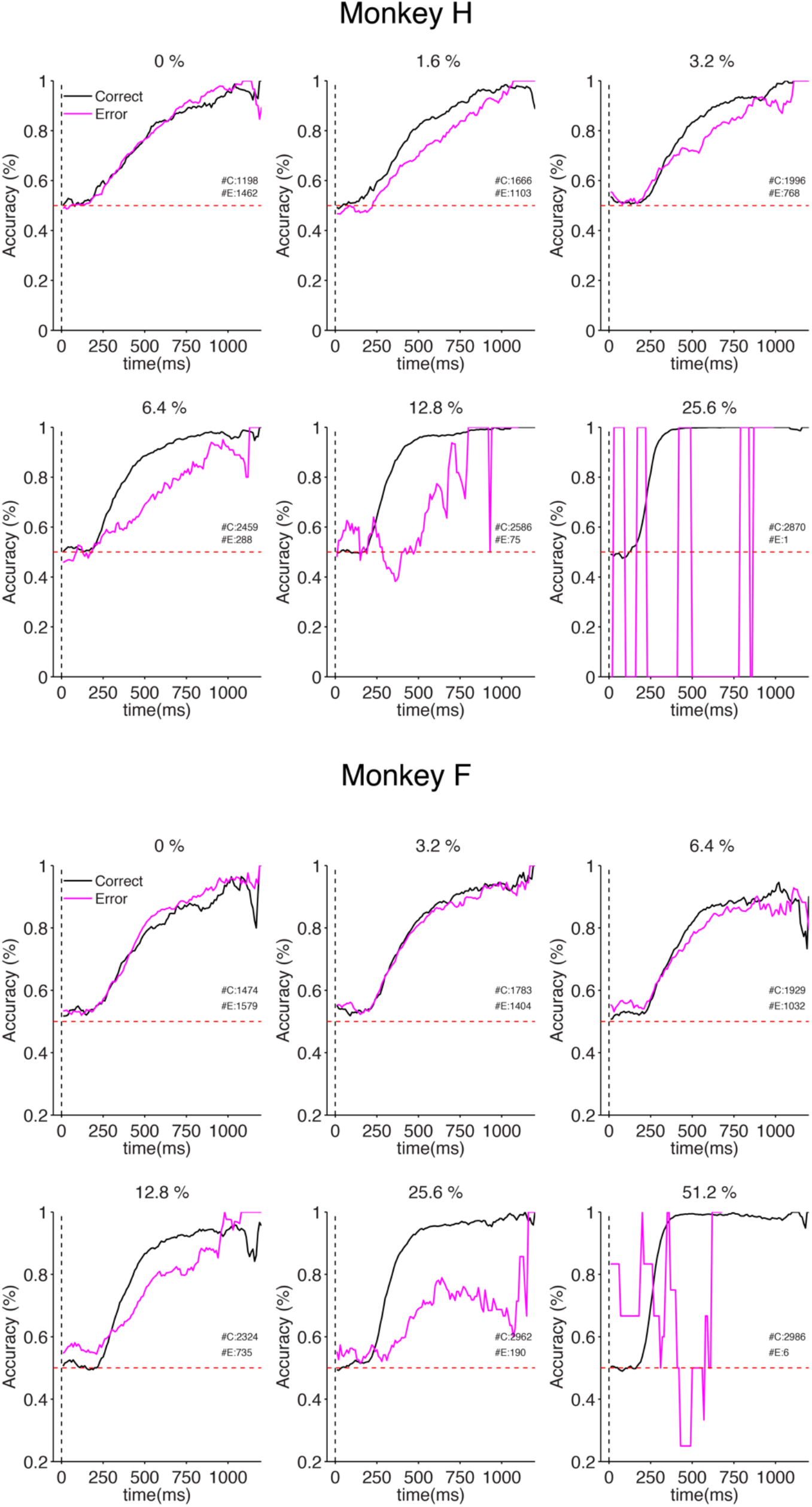
Choice prediction accuracy for correct and incorrect trials as a function of coherence. Choice prediction accuracy obtained from real-time readout for correct and incorrect trials for each level of coherence. Prediction accuracy during dots epoch for each coherence level is plotted for correct (black) and error (magenta) trials. Red dashed line corresponds to chance level. Insets show total number of Correct (C) and Error (E) trials used in the analysis. Data for monkey H and F are shown in top and bottom panels, respectively. Mean prediction accuracy for error trials after neural latency (180 ms after stimulus presentation) is outside (and lower than) the 95% CI for correct trials for 1.6%, 3.2%, 6.4%, 12.8% and 25.6% coherences for monkey H and for 12.8%, 25.6% and 51.2% coherences for monkey F - 1000 bootstrap iterations. Results for the highest coherence for each monkey should be interpreted carefully due to the extremely low number of error trials for these conditions resulting from excellent behavioral performance.

**Extended Data Figure 5.**
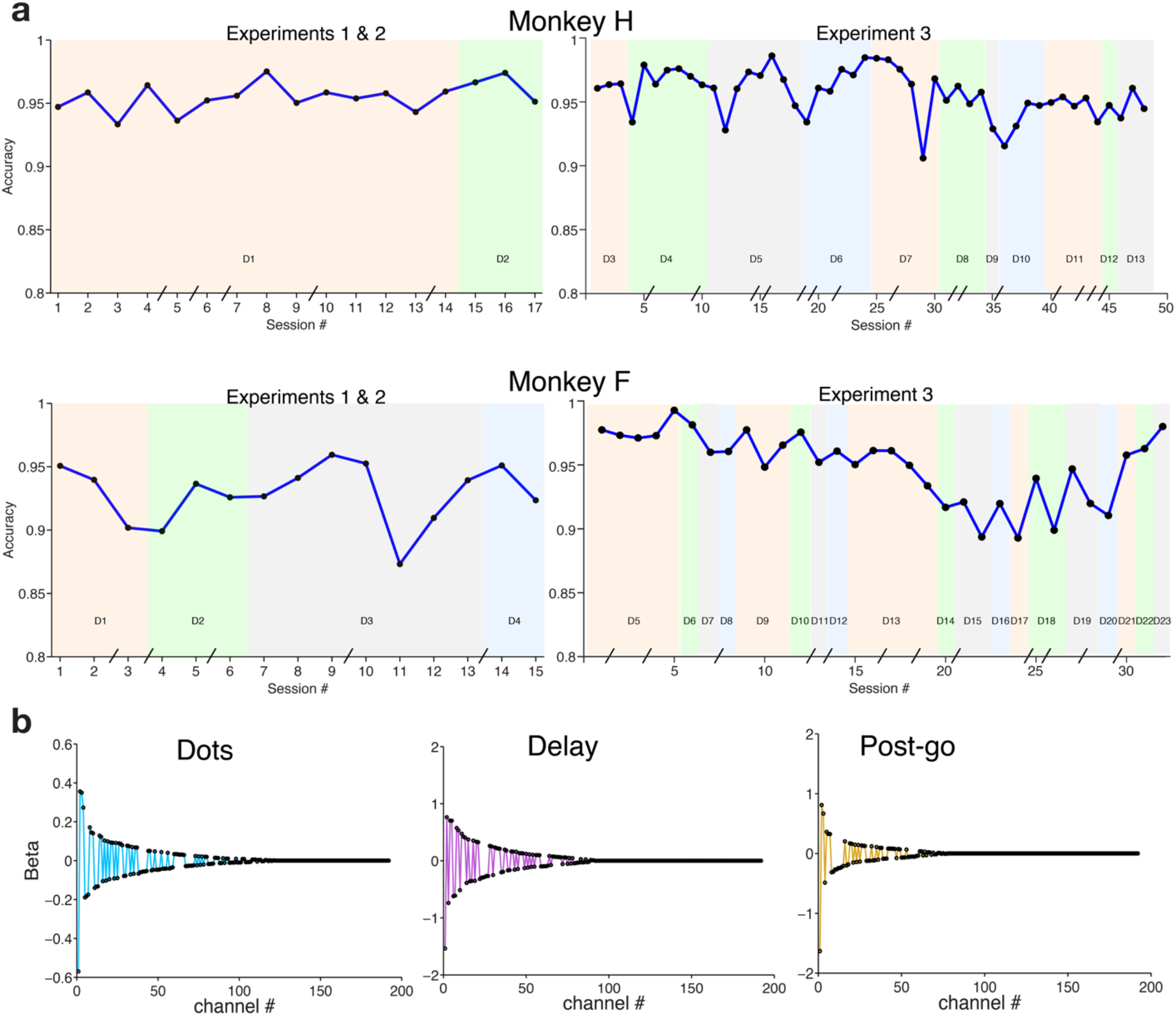
Real time decoding: performance reliability and decoder weights. **a) Decoding performance is stable across sessions.** Average prediction accuracy during the second half of the stimulus presentation (600-1200 ms) across all sessions for monkey H (top panels) and monkey F (bottom panels). D1-D23 denote different decoders (sets of beta weights) used for the recorded sessions. For monkey H the same decoder (D1) was used for the first 14 sessions. The breaks on the x-axis correspond to sessions that occurred on non-consecutive days. **b) Real time decoder beta weights.** Beta weights during the dots period (left panel) ranked by absolute magnitude for an example decoder used in real time experiments. Channels with no or little choice predictive activity during this period had their weights set to zero by using LASSO regularization to prevent over fitting. Delay period and Post go cue Beta weights are shown in the middle and right panels respectively.

**Extended Data Figure 6.**
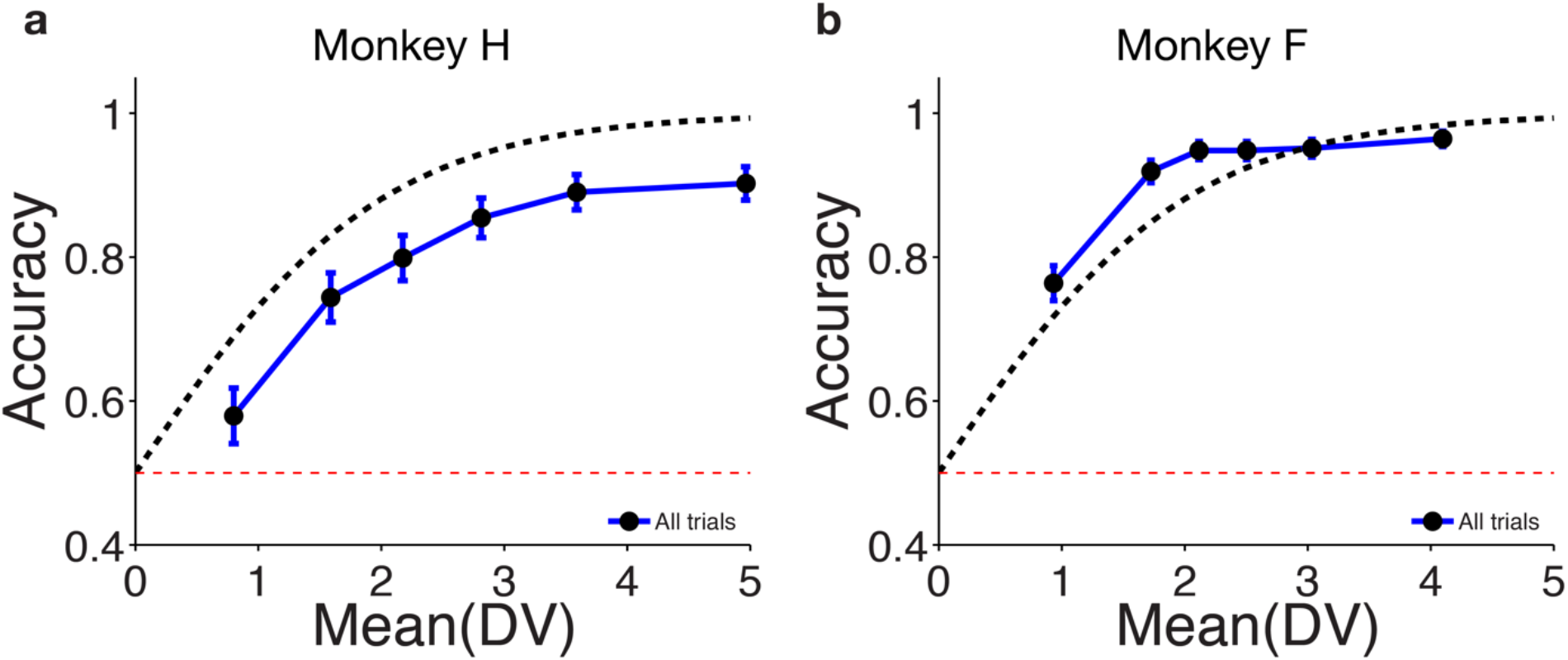
Prediction accuracy as a function of DV for CoM trials. **a) Choice prediction accuracy for all trials collected during the CoM detection experiment - monkey H.** Trials were split in 6 quantiles sorted by DV magnitude at termination. Prediction accuracy and median DV magnitude was calculated and plotted separately for each quantile (blue line with black markers). Blue error bars show standard error of the mean for a binomial distribution. Dashed black line shows predicted accuracy from log-odds equation and red dashed line shows chance level. Data from 985 CoM trials from monkey H. **b) Choice prediction accuracy for all trials collected during the CoM detection experiment - monkey F.** Same as **a)** for monkey F using 1727 CoM trials.

**Extended Data Figure 7.**
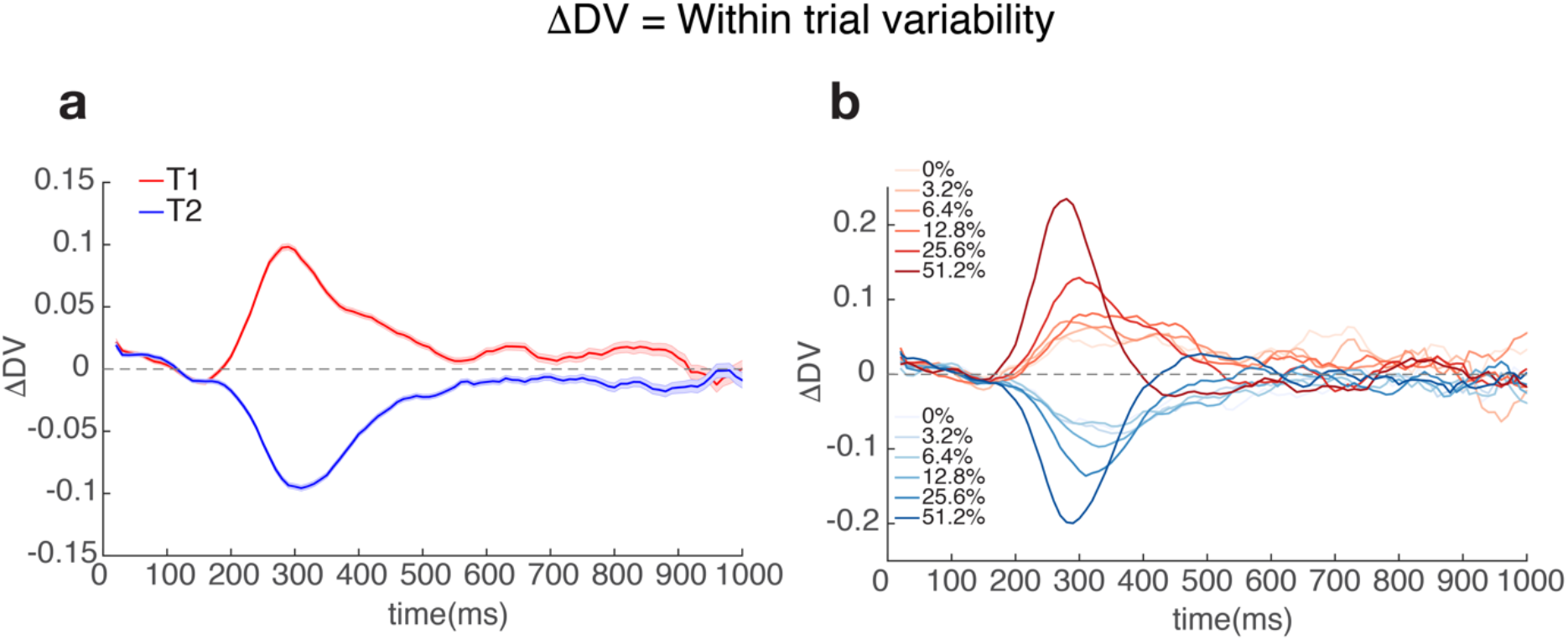
Within trial variability as a function of time, choice and stimulus coherence. **a) Average DV derivative as a function of time and choice - monkey F.** DV derivative was calculated for each trial as the difference between consecutive DV estimates spaced out by 10 ms. Traces show average DV derivative +/- s.e.m for right choices (red trace) and left choices (blue trace). **b) Average DV derivative as a function of time and signed coherence - monkey F.** Same data as in **a)** but with DV derivative averaged separately for each choice and motion coherence level (correct trials only). Right choices are plotted in red and left choices in blue as in **a)**. Darker traces correspond to stronger coherences.

**Extended Data Figure 8.**
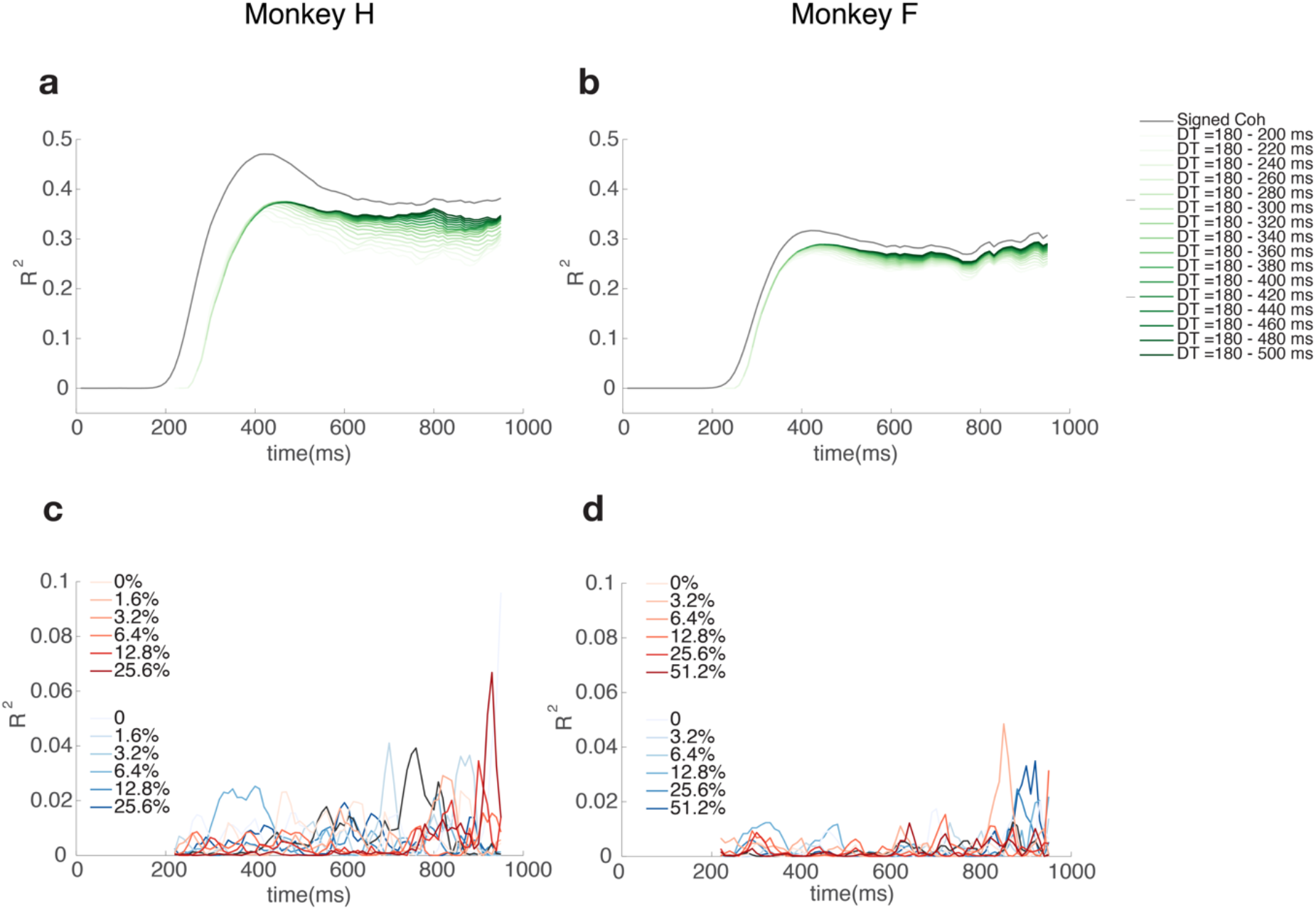
Correlation analysis between DV and stimulus motion energy. **a) Correlation between stimulus motion energy and decision variable - monkey H.** Proportion of variance explained when regressing decision variable as a function of signed stimulus coherence (grey trace) or motion energy (green traces). Each green trace corresponds to a separate regression between DV and average motion energy between a timepoint in the past (from −200 ms up to −500 ms) and −180 ms (the estimated neural response delay). Darker traces correspond to regressions in which the motion energy was averaged for a longer time window. Across all coherence levels motion energy and signed coherence explain a large fraction of DV variance. **b Correlation between stimulus motion energy and decision variable - monkey F**. Same as **a)** for monkey F. **c) Correlation between stimulus motion energy and decision variable within each level of signed coherence level - monkey H.** Proportion of variance explained when regressing DV for each time point and within each level of signed coherence as a function of the motion energy preceding it by 180 ms (the estimated neural response delay). Within each level of signed coherence, the DV fluctuations are not predicted by the motion energy traces **d) Correlation between stimulus motion energy and decision variable within each level of signed coherence level - monkey F.** Same as **c)** for monkey F.

## Methods

### Subjects

Our experiments were performed on two adult male macaque monkeys (*Macaca mulatta*) trained to perform a direction discrimination task with reaching movements of the arm as operant responses. These were the same subjects used in our previous study^10^, but with new experiments. All training, surgery, and recording procedures conformed to the National Institutes of Health Guide for the Care and Use of Laboratory Animals and were approved by Stanford University Animal Care and Use Committee.

### Apparatus

Monkeys sat in a custom-made primate chair (Stanford Machine Shop) in front of a video touchscreen, with their heads restrained using a surgical implant. The front plate of the chair could be opened, allowing the subjects to reach the touchscreen with the arm contralateral to the implanted hemisphere. The ispsilateral arm was gently restrained using a delrin tube and a cloth sling. Stimuli were shown on the video touchscreen (ELO Touchsystems 1939L), which was positioned approximately 35.5 cm away from the monkeys’ heads and allowed hand position to be tracked at 75Hz. Eye position was continuously tracked with an infrared eye tracker at 1kHz (EyeLink 1000, SR Research, Canada).

### Motion Discrimination Task

The task employed is a variation of the classical random dots motion discrimination task, in which the subject uses an arm movement as the operant response^10^ (Fig. 1a). We used a variable duration version of this task in which the duration of the stimulus presentation varied from trial to trial. There were two types of trials in our experiments: *open loop*, in which the stimulus duration was determined by the experimenter at the beginning of the trial and *closed loop*, in which the duration was contingent on a specific pattern of neural activity detected in real time (see Experiments 1-3). The subject was never cued on what type of trial it was on. For open-loop trials stimulus duration ranged from 500-1200 ms (median 670 ms) and was randomly chosen on each trial by sampling an exponential distribution. For closed-loop trials the possible values for duration ranged between 250-1200 ms and were determined on each trial either by the timepoint at which the termination conditions were met or a predetermined random duration sampled from the open loop distribution, whichever came first. All trials started with the onset of a fixation point (FP; 1.5 degree diameter) on the video touchscreen (Fig. 1a). To initiate the task, the monkey was required to maintain both eye and hand fixation within +/-3 degrees of the FP as long as it remained on the screen. Importantly, throughout the entire trial, the monkey was required to always maintain direct hand contact with the screen, otherwise the trial would be aborted.

After 300 ms of fixation, two targets (1.5 degree diameter) appeared on opposite sides of the FP (eccentricities between 10 and 17 degrees). After a 500 ms delay the random dot stimulus was presented for the durations mentioned above, after which it was removed from the screen. The monkey was asked to report the net direction of motion (0 or 180 degrees) by reaching to the target in the corresponding direction. The difficulty of the task was adjusted by changing the fraction of dots moving coherently in one direction (motion strength). After stimulus offset the monkey either entered a delay period during which it was required to withhold his response for 400-900 ms (on 30% of the open-loop trials) or was immediately presented the go cue (on 70% of the open-loop trials and all closed-loop trials). The go cue was then signaled by the offset of the FP at which point the monkey was free to gaze anywhere and report his decision with his arm by reaching one of the two targets. Although gaze was monitored, reward acquisition depended solely on reaching to the correct target. Finally, for a response to be considered valid, the monkey was required to hold its hand position within +/-4 degrees of the center of the target for 200 ms. The monkey was then rewarded with a drop of juice for correct choices and given a timeout (2-4 seconds) for incorrect ones. Zero coherence trials were rewarded randomly with a probability of 0.5 since there was no correct response on these trials. The motion discrimination task was run on an Apple Mac Pro running Mac OS.

### Random dots stimuli

The stimuli used in our psychophysical experiment were random dot kinematograms (RDK) generated using MATLAB and Psychophysics Toolbox. The details for generating the random dots stimuli have been described previously^10^. However, to allow for closed loop experiments 1 and 2 (see below) we introduced a modification to be able to terminate the dot presentations early if needed. The stimulus code was designed to precompute a sequence of kinematograms that contain both random and moving dots. The sequence was then presented ballistically with no need to continuously compute the content of each frame. Our modification allowed for DV values to be received asynchronously from the real-time decoder and evaluated during the dots presentation. If the DV criteria defined by the particular experiment were met, the dot presentation could then be terminated without the remaining frames being shown. For the experiment in which an additional pulse of motion energy was injected (closed loop experiment 3, see below), we arranged for two sequences of kinematograms to be precomputed before presentation: one without the pulse, the other for the 200 ms pulse itself. Contingent on the evolution of DV values, the stimulus could then be rapidly switched from the standard sequence to the pulse sequence.

For both monkeys, the motion strength could take one of 6 possible values within a set, but the sets were slightly different between subjects: [0%, 1.6%, 3.2%, 6.4%, 12.8%, 25.6%] for monkey H and [0%, 3.2%, 6.4%, 12.8%, 25.6%, 51.2%] for monkey F. The top coherence (51.2%) was dropped and a very low coherence (1.6%) was introduced for monkey H, due to its superior discrimination ability. The direction and coherence of the motion were randomly assigned on each trial by sampling from a uniform distribution with replacement. For zero-coherence stimuli all dots were displaced randomly but, due to the stochasticity of that process, one obtains non-zero net motion toward the targets over a small number of frames.

### Behavioral Training

Both monkeys had been extensively trained on fixed and variable duration versions of the motion discrimination task using an arm reach movement as the operant response prior to the current study^10^. A few training sessions (all open-loop trials) were used to get the subject accustomed to the new task timing (0.5-1.2 s stimuli and no delay on 70% of the trials). Real time decoding sessions only started when psychophysical performance was stable.

### Behavioral Analysis

Psychophysical performance was assessed in two ways: by describing the percentage of correct choices as a function of (unsigned) stimulus coherence and by describing the percentage of rightward choices as a function of signed stimulus coherence.

The percentage of correct choices as a function of motion strength (stimulus coherence) was fit by a cumulative Weibull distribution function:

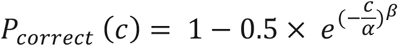

where *P_correct_*is probability correct, *c* is motion strength, *α* is the psychophysical threshold (the value of *c* that corresponds to ∼82% correct responses), and *β* is a parameter that controls the shape of the function, especially its steepness.

The proportion of rightward choices, *P_right_,* as a function of motion strength and direction was fit by a logistic regression:

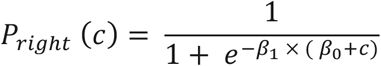

where *c* is motion strength, *β_1_* is the slope parameter and *−β_0_* is the motion strength corresponding to the indifference point. This value was used to assess the monkey’s behavioral bias on each session.

### Electrophysiological recordings

Two multielectrode arrays (Blackrock Microsystems, Utah) with 96 electrodes each (1mm long platinum-iridium electrodes, 0.4 mm spacing, impedance average of approximately 400 KOhm) were implanted in primary motor and dorsal premotor cortex of each monkey (Figure 1c). The methods for determining the array placement were described in our previous study^10^. For monkey F, the M1 array became unusable between the end of the previous study and the start of the current study. Due to lack of neural signal from the M1 array, only the PMd array was used for this animal. Continuous neural data were acquired and saved to disk from each channel (sampling rate 30 kHz) and thresholded at −4.5 RMS using the Cerebus recording system (Blackrock Microsystems, Utah) and two separate PCs (one for each array) running Windows 8. Waveforms corresponding to threshold crossings were not sorted and each channel could contain one or more unit(s). Sorting waveforms would require a significant lead-up time before the beginning of the experiment and could negatively affect the ability to combine data and use decoders across days (see below, Decoder training). Since units were not isolated within each channel our resulting units were most likely multi-unit clusters. Any extremely noisy channels were deactivated at the beginning of a session, and all other channels were used in this study. Using only multi-units yielded comparable prediction accuracy (Extended Data Figure 3) to our previous study^10^ in which both single and multi-unit data was used.

### Datasets

Data were collected in two sets of experiments. In the first set of experiments we performed Closed Loop Experiments 1 and 2 (see below). For this set, for each monkey we analyzed all datasets that met two behavioral inclusion criteria: 1) over 500 trials and 2) a behavioral bias (|β_0_|) under 4%, as determined by a logistic regression fit (see above). These criteria were imposed to ensure that we have a sizeable number of trials per condition (6 coherence x 2 directions = 12 conditions) and that the behavior of the monkey is virtually unbiased, such that both neural and behavioral results are more easily interpretable. These criteria resulted in a selection of 17/15 sessions for a total of 16468/15826 trials for monkey H/F, respectively.

In the second set of experiments we performed Closed Loop Experiment 3. This set of experiments was performed later, on separate sessions, but using the same two subjects, arrays and decoding techniques as the first set. In this set of experiments, we analyzed all datasets with over 550 trials. For all experiments in monkey H, PMD and M1 were recorded simultaneously.

### Decoder training

We chose to use a logistic regression classifier based on our previous results showing excellent offline prediction accuracy in variable duration tasks^10^ and because of the direct probabilistic interpretation of its output. Our decision variable (DV) was defined as the log odds ratio of observing a particular behavioral choice (*T_1_*or *T_2_*) given the population response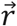:

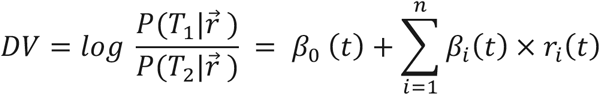

Where *r_i_*(*t*) are the z-scored summed spike counts for each neuron and time window, *β_0_* is an intercept term and *β_i_(t)* are the classifier weights (one for each unit and epoch). Data from all electrodes with valid waveforms were combined.

For simplicity, we decided to use only 3 different decoders for an entire trial (Fig. 1a), instead of a different one for each 50 ms time window in the trial^10^. We applied the first decoder from fixation up to and including the dots period, the second for the delay period and the third for the post go cue period. After extensive offline tests on a few sessions the precise epochs for classifier training were defined as the following:

- Dots epoch: [150, 1000] ms aligned to dots onset;
- Delay epoch: [250, 350] ms aligned to dots offset;
- Post-go cue epoch: [200,400] ms aligned to go cue;

LASSO regularization was applied to prevent over-fitting when calculating each set of beta weights. A Lambda parameter constraining the L1 norm of the Beta vectors was calculated separately for each of the 3 decoders using 10-fold cross validation on the corresponding time epochs listed above. For each decoder the Lambda value with minimum cross-validation error was chosen. Extended Data Figure 5b shows beta weights for an example set of 3 decoders for monkey H sorted by epoch and ranked by magnitude. Positive weights correspond to rightward preferring channels while negative weights correspond to leftward preferring channels. LASSO regularization sets weights of channels with little or no predictive activity to zero.

The linear classifier was determined offline using recently collected data (from real-time experiments). All 50 ms samples of neural data during the selected period (above) for each epoch were used to train the classifier. The classifier was trained on 90% of the trials and tested on 10% of the trials using 10-fold cross-validation. The weights from one of the cross-validation folds were then used in the upcoming real-time experiments. Decisions to train new decoders were based on experimenter judgment in attempts to optimize performance: if a substantial decrease in real-time decoding performance and/or an increase in the DV offset at baseline was observed, a new classifier was trained and used in the following session. New classifiers were typically used every 5 sessions, but some proved to be stable over up to 14 sessions (Extended Data Figure 5a).

### Real time decoding

An essential requirement to compute a real-time read-out of neural activity is the ability to continuously and (nearly) instantaneously access and perform computations on the neural activity being recorded. To accomplish this, the spikes for each channel were temporally smoothed using a causal half-Gaussian kernel with 50 ms standard deviation (to mitigate spurious Poisson fluctuations) and summed for the most recent 50 ms. These smooth spike counts were then stored in a 192×1 (96×1 for monkey F) vector of neural activity and z-scored individually for each channel, using previously calculated µ (mean) and σ (standard deviation) vectors. Z-scoring neural activity was crucial to ensure a reliable and stable real-time readout by preventing the highest firing channels from dominating it. Finally, the z-scored neural activity was projected onto a previously calculated linear decoder (a set of β weights, one for each channel) to obtain our linear readout of internal decision state: a real time decision variable (DV)^1^.

The value of the DV was updated every 10 ms, reflecting the neural activity of the preceding 50 ms. Because we used a half-gaussian kernel, data preceding the 50 ms window also influenced our DV estimate (with more recent spikes carrying more weight). 95% of the data contributing to the spike counts was limited to the last 100 ms (i.e an additional 50 ms in the past to each 50 ms window). The DV value and its history on a single trial could then be used (if desired) to impose conditions for termination of the random dots stimulus (experiments 1 and 2) or presentation of a motion pulse (experiment 3), effectively closing the loop on the experiment.

While the β weights were not updated online (during the course of one experiment), the µ and σ vectors for each epoch were learned continuously during the course of the experiment, due to changing recording conditions and signals from day to day. The µ and σ vectors were initialized at the beginning of the session using the values calculated offline when training the most recent decoder. Once the session started, the initial µ and σ vectors were blended with online calculated values for the first 25 trials, using a blending factor α:

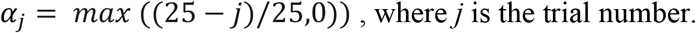

For trial *j*, sample number t and for a given epoch in trial, the µ and σ vectors were defined as a weighted mixture between the initial values µ_initial_(epoch) and σinitial(epoch) and the estimate of the current session’s values µ_current_(t,epoch) and σ_current_(t, epoch):

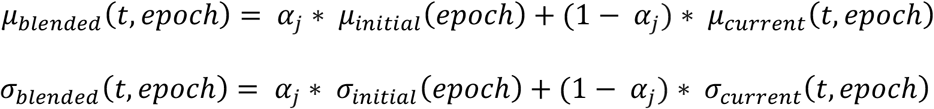

After the first 25 trials α was set to zero which implies the µ and σ vectors kept being continuously updated throughout the session but were no longer blended with values from the previous days. The update rule for µ_current_(t, epoch) was:

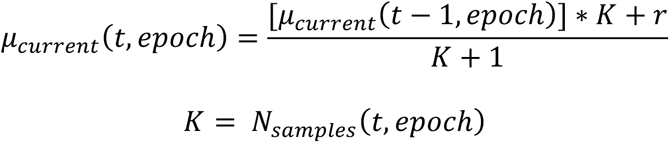

where r is the most recently sampled vector of spike counts and *K* is the current number of samples of spike count vectors obtained so far for this particular epoch.

The update rule for σ_current_(t, epoch) was:

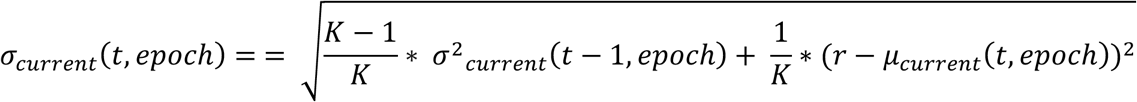

After updating the µ_current_(t, epoch) and σ_current_(t, epoch) vectors, the number of samples for the corresponding epoch was also updated:

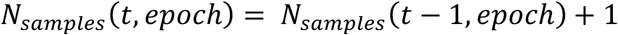

Importantly, even though we had only 3 different decoders (Fig. 1b) we effectively used 5 different epochs: Fixation, Targets, Dots, Delay and Post Go-Cue. The Dots decoder was also used in the Fixation and Targets epochs but because average firing rates are different between these, different µ and σ vectors had to be used. Every 50 ms sample of neural data for a given epoch was used to update the corresponding µ and σ vectors as described above. We let the µ and σ vectors converge for ∼200-300 trials, in the beginning of each experimental session, before starting any closed loop experiments. One way to check for this convergence was to monitor the DV offset: the average DV value for the first 150 ms of the Dots epoch. Since we verified through offline analyses that no systematic pre-planning activity towards one of the two targets was present in PMd or M1 during this time window, we expected the DV offset to be on average ∼0.

Using a single decoder for an entire epoch was far more efficient to implement than using a different decoder for each time point (as it reduced the number of µ and σ vectors that had to be learned online) and as demonstrated in our previous study^10^ a single classifier for an entire epoch was almost as predictive as multiple classifiers trained on different timepoints of the same epoch^10^. Because choice modulation in PMd/M1 changes dramatically around the peri-movement period a single decoder for an entire trial was not feasible.

In the end, our method yielded a reliable real-time decision state read out and required only ∼18% (15%) of trials in a session for calculating the values of µ and σ for monkey H (F), leaving the remainder available for imposing neurally contingent conditions in closed loop. The real time decoder was run on two separate PCs (server and client) using the Simulink Real-Time/xPC platform (Mathworks, Massachussetts).

### Closed Loop Experiments

#### Experiment 1: Virtual Boundaries

On each trial we set a virtual threshold, or boundary (B), for the magnitude of the DV during the dots epoch. If the DV on the current trial reached B or −B ± tolerance, the dots presentation was terminated and the monkey asked to report its decision. If the bound was not reached on a given trial, stimulus presentation continued to a preset duration for that trial which was randomly sampled from an exponential distribution ranging from 500-1200 ms. Closed loop trials for which the boundary was not reached were effectively indistinguishable from open loop trials.

Typically, 5 values for boundaries spanning 0.5 to 5 (DV units) were used every session and one of them was randomly assigned on each trial (uniform distribution). The tolerance used was ± 0.25 DV units. We imposed a minimum duration for all trials to avoid spurious bound crossings, which could be problematic for low bound values in particular. In all sessions the minimum duration was 250 ms, a conservative estimate of the latency for choice related signals driven by the visual stimulus to appear in PMd and M1.

After the minimum stimulus duration was reached, the DV was assessed every 10 ms to determine whether it fell within ± 0.25 DV units of the bound chosen for the current trial (B or −B). If so, the stimulus was terminated within 34 ms of the boundary being met (see Estimated latency for real time closed loop setup). If the bound for the particular trial was not reached, the presentation continued up to the maximum stimulus duration selected for that trial which had been obtained by randomly sampling from an exponential distribution: 500-1200 ms (median 670 ms).

Finally, closed loop trials were randomly interleaved with open loop trials in which no DV-dependent termination condition was imposed. The motivation for interleaving closed loop and open loop trials was to make it extremely hard for the monkey to learn that accelerating the dynamics of choice related signals^10^ (potentially by recruiting more choice related neurons or increasing their modulation) and thus hitting bounds sooner could potentially increase its reward rate. Not accounting for this possibility could lead to an undesirable change in the monkey’s strategy during the course of the closed loop experiments, which could become problematic when combining data across days.

#### Experiment 2: CoM detection

Under our logistic regression framework, the signature of a putative CoM is a sign change of the decision variable. Since these sign changes could happen at any time during the trial, capturing them required not only monitoring the most recent state of the DV, but its history throughout the trial. Because there was noise in our DV estimation and DVs usually started close to 0 at the beginning of the trial we imposed selection criteria to detect likely CoMs based on the neural data. A necessary feature for all potential CoMs was a zero crossing in the sign of the DV: change of DV sign from negative to positive reflected a change in the likelihood of a rightward decision from less than 50% to greater than 50%, and vice versa for the opposite change in sign. To eliminate zero crossings resulting solely from measurement noise, we imposed four additional criteria:

- Minimum DV value after zero crossing;
- Minimum DV value with opposite sign before zero crossing;
- Minimum duration of DV sign stability after zero crossing;
- Minimum duration of DV sign stability before zero crossing;

The minimum DV values before and after zero crossing were symmetrical for most sessions, as were the periods of minimum duration of DV sign stability (negative or positive values for all time points). If a zero crossing was detected and all four criteria were met, the stimulus presentation was interrupted and the animal was virtually immediately (within 34 ms or less, see Estimated latency for real time closed loop setup) prompted to report a decision. The exact parameters used for each session can be found in Supplementary Information Table 3.

By sweeping the parameter space we could test zero crossings that differed in magnitude and stability. Analogously to the virtual boundary experiment, if the minimums were not met and a CoM thus not detected, the stimulus presentation continued uninterrupted for a random duration ranging from 500-1200 ms, selected prior to the start of the trial. A minimum stimulus duration of 250 ms was also in place.

Because putative CoMs are quite rare^1^, in the first set of experiments we devoted 70% of the closed loop trials to detect them leaving the remaining 30% as virtual boundary trials. The exact fraction of trials with CoM depends dramatically on how we parameterize them. The longer the minimum periods of consistent sign and the higher the minimum DV value in the initial commitment stage, the rarer they become. Running both experiments on the same sessions ensured that the mapping from DV to choice likelihood was held during the CoM experiments and provided the most faithful indirect validation of initial commitment we could obtain.

#### Experiment 3: Motion pulse perturbation

In this experiment, motion pulses were introduced on some trials with motion coherences near or below psychophysical threshold. No motion pulses were presented for suprathreshold coherences based on the results of a pilot experiment (not shown) in which pulses presented at suprathreshold coherences were more perceptually salient and led to changes in the animals’ strategy. As in Experiment 1, on each trial we set a virtual boundary (B) for the magnitude of the DV during the dots epoch. In this experiment, 100% of trials with dots coherence at or near psychophysical threshold were treated as closed loop trials (this corresponds to trials with maximum unsigned coherence of 6.4% for monkey H and 12.8% for monkey F; psychophysical thresholds were measured using the Weibull function described above in “Behavioral Analysis”). Low-coherence trials in which the boundary was not reached (per the criteria below) and trials with suprathreshold dots coherences were all effectively open loop.

If the DV on a closed loop trial reached B or −B ± tolerance (±0.25 DV units), after a minimum stimulus duration of 50 ms, a 200-ms motion pulse was presented, followed immediately by termination of the visual stimulus and presentation of the cue for the monkey to report its decision.

If not, dots presentation continued for a pre-assigned duration drawn randomly from an exponential distribution of 500-1200 ms. Four integer values for boundaries (spanning 1 to 4 DV units) were used every session, and one of them was randomly assigned on each trial (uniform distribution).

Motion pulses were 200-ms periods of additional dots stimulus presentation with small additive average coherence (±2% or 4.5% from the initial dots coherence on the same trial for monkey H and F, respectively, where positive coherence values indicate rightward motion); thus pulses effectively randomly added either a small amount of rightward or leftward motion evidence to the stimulus. Pulse strength was calibrated in pilot experiments, in which we converged upon coherence shifts that slightly but significantly biased each animal’s behavior, without being overtly perceptually salient (biases were measured using the logistic regression on rightward choice described above in “Behavioral Analysis”). Animals were rewarded for correct reaches in the direction of the coherence of the initial dots stimulus (randomly assigned on 0% coherence trials), regardless of the pulse direction.

#### Estimated latency for real time closed loop setup

To validate our setup, we measured the latency between a neural condition being met and the corresponding task change being implemented. We tested this latency by generating simulated DV steps in the same model used to detect when DV triggering conditions were met in the real experiments. We used these simulated steps to trigger the onset of a bright light on the touchscreen in front of a photodetector, again within the same code used to run the task and generate the stimuli in the real experiments. We then passed both the simulated DV and the photodetector output signals into an oscilloscope, triggered the display on the “DV” steps, and manually measured the delay to onset of the bright dot. Almost all measured delays were within 2 frames, or 26 ms.

#### Estimated trial count savings for real time closed loop setup

The real time setup allowed for extremely precise experimental control over which DV values or DV history to use to trigger a modification in the task (stimulus termination or pulse). However, it could be argued that given enough data, similar trials would have been captured simply by either terminating the stimulus (as in experiments 1 and 2) or presenting the pulse (as in experiment 3) at a random point in the trial and then back sorting them offline (by DV value or history by after the data is collected).

To estimate how much more trial-count efficient it was to use our real time setup compared to offline back-sorting trials where the stimulus was presented for a random duration, we used the CoM experiment as a case study given how rare change of mind events are.

For simplicity, we focused on sessions 1, 2 and 3 from Monkey F, which all have the same (and intermediate) CoM requirements (Supplementary Information Table 3). We started by calculating the yield from the real time experiment in closed loop as the ratio between detected CoM trials and trials in which CoMs were checked (i.e. all closed loop trials in which the stimulus could be terminated if the conditions dictated by the CoM parameters were met, Supplementary Information Table 3):

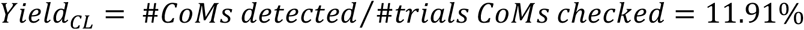

To calculate the yield for offline back-sorting trials we used the open loop trials in the same sessions, which were terminated after a random stimulus duration. Importantly, the stimulus duration on these open loop trials was sampled from the same distribution as for the closed loop trials in which CoMs were checked, which allows for a fair yield comparison. We calculated the yield from offline back-sorting as the ratio between the number of trials that would have met all the criteria for CoMs for the same session and the total number of open loop trials:

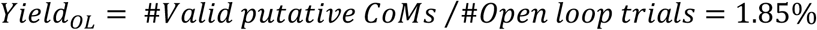

Since the goal would be to probe the new choice preference shortly after the zero crossing (putative change of mind), not many hundreds of ms later, we only considered CoM trials that were (closed loop) or that would have been (open loop) terminated within 150 ms of the zero crossing. This cutoff value corresponded to the 82^nd^ percentile of post zero crossing durations for the closed loop trials analysed in these sessions.

In this analysis Yield_CL_was 6.43 times higher than Yield_OL_.This result implies that had we not used a real time setup in closed loop we would have had to collect 6.43 times the number of trials (and thus sessions) to obtain the same number of events. This would in turn mean collecting around 100 sessions/monkey just for experiments 1 and 2 (assuming the same 30%/70% trial split used in the real time experiments), rendering this experiment practically unfeasible.

### Neural Data Analysis

#### DV variability

Within trial variability was computed by first calculating the difference between consecutive DV values (estimated every 10 ms) for every trial in the datasets collected for experiments 1 and 2 (open and closed loop). This step yielded a ΔDV trace for each trial aligned to dots onset. For each trial these traces were computed only up to the offset of the stimulus and did not include any delay or post go-cue DV data. The ΔDV traces were then sorted and averaged for each choice (Fig. 5a, Extended Data Fig.7a) or each signed coherence level (Fig. 5b, Extended Data Fig.7b). Longer trials are increasingly rare due to the shape of our stimulus duration distribution but this asymmetry does not influence the interpretation of the time course of average ΔDV as this metric only captures *within* trial variability and not *across* trial variance.

#### DV and motion energy correlation

Motion energy (ME) was calculated for each trial in the datasets collected for experiments 1 and 2 (open and closed loop) by convolving the positions of the dots in the stimulus with spatio-temporal filters as previously described^8^. The ME trace obtained for each trial captures the strength of the stimulus at every timepoint during the stimulus presentation. To evaluate the effect of motion energy on DV we performed a linear regression of single trial DV traces on single trial ME traces. From experiments 1 and 3 we determined that due to neural latencies a stimulus fluctuation can only have effect on the decoded DV ∼180 ms later. For this reason, the regression was always performed between DV(t) and ME(t-180ms) or earlier. For Extended Data Fig. 8a-b each green trace corresponds to a different way to estimate the motion energy that might affect DV at time t. For the lightest trace and for every timepoint t ME was averaged between (t-180ms) and (t-200ms) for every trial and used to regress against DV(t). A separate regression was performed for each timepoint and the resulting variance explained was plotted. The same process then was repeated for every other green trace by progressively increasing the averaging window for ME in 20 ms increments from (t-180, t-200) ms to (t-180, t-500) ms. As a control the DV was also regressed against signed coherence for each trial (Extended Data Fig. 8a-b grey traces). This analysis was used to assess how much of the DV variance *across* coherences is explained by motion energy or signed coherence as a function of time. To assess how much DV variance *within* each coherence level could be explained by the motion energy of the stimulus we first sorted the DV traces for each signed coherence level. For each signed coherence level and each timepoint we regressed DV(t) against ME(t-180ms) for the corresponding trials and calculated the variance explained (Extended Data Fig. 8c-d).

#### CoM regularities

To test whether the effects of coherence on the number of CoMs were statistically significant we used a bootstrap method to generate 1000 distributions of CoM events with the corresponding coherences by sampling with replacement from the distribution of captured events for each subject separately. For each subject each distribution had the same number of observations as those captured in experiment 2: 985 for monkey H and 1727 for monkey F. For each randomly sampled distribution the number of CoMs for each coherence level was counted. The resulting counts were then regressed against the coherence level they belonged to. CoM count was highly and negatively correlated with coherence for both subjects (p<0.001).

To test whether the effect of directionality of CoMs (corrective vs erroneous) was statistically significant we followed a similar bootstrapping procedure and generated 1000 distributions of CoM events (excluding 0% trials) with the corresponding directionality. For each randomly sampled distribution the number of CoMs for each direction was counted. The difference between the medians counts of corrective and erroneous CoMs was tested by performing a one-sided Wilcoxon rank sum test (p<0.001) testing the hypothesis than corrective counts were higher than erroneous counts.

#### Pulse Effects

To quantify the overall behavioral effect of the pulses, we performed the following logistic regression on the probability of a rightward choice:

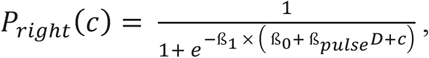

where *c* is motion strength, *β_1_* is the slope parameter, *D* is the pulse direction, and *−β_0_* is the motion strength corresponding to the indifference point.

To determine the effect of the pulse on the evolving DV, we first estimated the minimum latency for the visual stimulus information to influence the DV by calculating the first time of significant divergence of rightward vs. leftward DV traces during dots presentation on open-loop trials with stimuli of maximal motion strength (±25.6%, 51.2% coherence for monkey H, F), assessed using a two-sample t-test with correction for a false discovery rate of 0.05^34^. We refer to this estimate as the “evidence representation latency” (ERL; 170 ms, 180 ms for monkey H, F). We then measured the evolution of the DV after pulse presentation by calculating the difference between the empirically observed DV at each time point *t* and the DV at the “pulse evidence representation latency” (PERL, or time of pulse onset plus the ERL):

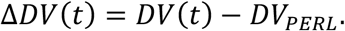

To quantify the pulse effect on DV on single trials, we calculated the slope (using linear regression) of ΔDV over the time period beginning with the PERL and ending at either the median go-RT for the animal or 150 ms prior to movement initiation on that trial, whichever came first, to minimize the contribution of directly movement-related activity to our analysis of the evolving choice^10^. We checked to ensure the results did not depend critically on these specific endpoint criteria by sweeping a range of endpoints for the ΔDV calculation: every 10 ms from go cue onset to 150 ms after the go cue. The exact endpoint used did not affect the results of the following analyses (quantitative results held for all endpoints from the time of the go cue through 150 ms post-go-cue for monkey H, and for endpoints beginning 120 ms post-go-cue for monkey F).

To quantify the neural pulse effect at each DV boundary (as in Fig. 5e), we fit the following linear regression on the ΔDV slope (calculated as described in the previous paragraph) separately for each DV boundary:

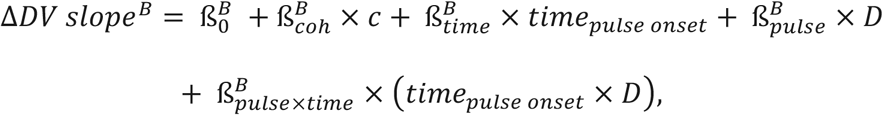

where *B* is the pulse-triggering DV boundary (unsigned), and compared 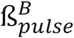 values. (The full resulting sets of regression coefficients fit to ΔDV slope can be found in Supplementary Information Table 1.)

Similarly, to quantify the behavioral pulse effect at each DV boundary (as in Fig. 5f), we fit the following logistic regression on the probability of a rightward choice separately for each DV boundary:

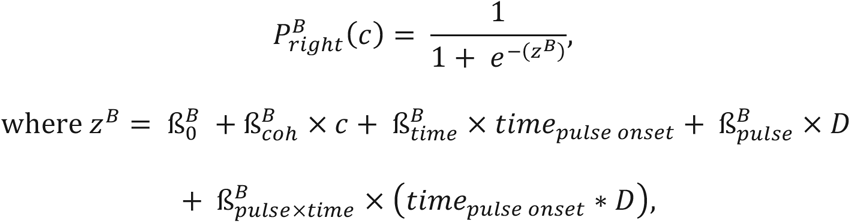

where *B* is again the pulse-triggering DV boundary (unsigned), and compared 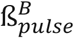 values. (The full resulting sets of regression coefficients fit to choice can be found in Supplementary Information Table 2.)

### Data availability

All data and analyses generated during the current study are available from the corresponding authors upon reasonable request.

### Code availability

The analysis code was developed in Matlab (Mathworks) and is available from the corresponding authors upon reasonable request.

## End notes

**Supplementary Information** is linked to the online version of the paper.

## Acknowledgments

We thank all other members of the Newsome and Shenoy labs at Stanford University for comments on the methods and results throughout the execution of the project. D.P. was supported by the Champalimaud Foundation, Portugal, and Howard Hughes Medical Institute. J.R.V. was supported by Stanford MSTP NIH training grant 4T32GM007365. R.K. was supported by Simons Collaboration on the Global Brain grant 542997, Pew Scholarship in Biomedical Sciences, National Institutes of Health Award R01MH109180, and a McKnight Scholars Award. J.C.K. was supported by NSF graduate research fellowship. P.N. was supported by NIDCD award R01DC014034. C.C. was supported by K99NS092972 and 4R00NS092972-03 award from the NINDS and supported as a research specialist by the Howard Hughes Medical Institute. J.B. and S.F. were supported by the Howard Hughes Medical Institute. K.V.S. was supported by the following awards: NIH Director’s Pioneer Award 8DP1HD075623, DARPA-BTO ‘‘NeuroFAST’’ award W911NF-14-2-0013, the Simons Foundation Collaboration on the Global Brain awards 325380 and 543045, and ONR award N000141812158. W.T.N. and K.V.S. were supported by the Howard Hughes Medical Institute.”

## Author Contributions

D.P., J.R.V., R.K., S.F., K.V.S. and W.T.N. designed the experiments. D.P., J.R.V. and S.F. trained the animals and collected the data. D.P., J.R.V. and W.T.N. wrote initial draft of the paper. S.I.R, D.P. and R.K. performed the surgical procedures. D.P., J.C.K., P.N., C.C. and J.B. implemented the real-time decoding setup. D.P., R.K. and C.C. designed, and D.P. and J.R.V implemented, the decoder training algorithm to obtain the decoder weights and normalization matrices. D.P. and J.R.V. analysed the data. All authors contributed analytical insights and commented on statistical tests, discussed the results and implications, and contributed extensively to the multiple subsequent drafts of the paper.

## Author Information

The authors declare no competing interests. Correspondence and requests for materials should be addressed to D.P., J.R.V., or W.T.N.

## References

1 Kiani, R., Cueva, C. J., Reppas, J. B. & Newsome, W. T. Dynamics of neural population responses in prefrontal cortex indicate changes of mind on single trials. Curr Biol 24, 1542–1547, doi:10.1016/j.cub.2014.05.049 (2014).

2 Kaufman, M. T., Churchland, M. M., Ryu, S. I. & Shenoy, K. V. Vacillation, indecision and hesitation in moment-by-moment decoding of monkey motor cortex. Elife 4, e04677, doi:10.7554/eLife.04677 (2015).

3 Bollimunta, A., Totten, D. & Ditterich, J. Neural Dynamics of Choice: Single-Trial Analysis of Decision-Related Activity in Parietal Cortex. The Journal of Neuroscience 32, 12684 (2012).

4 van den Berg, R. et al. A common mechanism underlies changes of mind about decisions and confidence. eLife 5, e12192, doi:10.7554/eLife.12192 (2016).

5 Lemus, L. et al. Neural correlates of a postponed decision report. Proceedings of the National Academy of Sciences 104, 17174, doi:10.1073/pnas.0707961104 (2007).

6 Rich, E. L. & Wallis, J. D. Decoding subjective decisions from orbitofrontal cortex. Nat Neurosci 19, 973–980, doi:10.1038/nn.4320 (2016).

7 Resulaj, A., Kiani, R., Wolpert, D. M. & Shadlen, M. N. Changes of mind in decision-making. Nature 461, 263, doi:10.1038/nature08275 https://www.nature.com/articles/nature08275#supplementary-information (2009).

8 Kiani, R., Hanks, T. D. & Shadlen, M. N. Bounded integration in parietal cortex underlies decisions even when viewing duration is dictated by the environment. J Neurosci 28, 3017–3029, doi:10.1523/JNEUROSCI.4761-07.2008 (2008).

9 Britten, K. H., Shadlen, M. N., Newsome, W. T. & Movshon, J. A. The analysis of visual motion: a comparison of neuronal and psychophysical performance. J. Neurosci. 12, 4745–4765 (1992).

10 Peixoto, D. et al. Population dynamics of choice representation in dorsal premotor and primary motor cortex. bioRxiv (2018).

11 Shadlen, M. N. & Newsome, W. T. Neural Basis of a Perceptual Decision in the Parietal Cortex (Area LIP) of the Rhesus Monkey. J Neurophysiol 86, 1916–1936 (2001).

12 Ratcliff, R. & McKoon, G. The Diffusion Decision Model: Theory and Data for Two-Choice Decision Tasks. Neural Computation 20, 873–922, doi:10.1162/neco.2008.12-06- 420 (2007).

13 Mazurek, M. E. A Role for Neural Integrators in Perceptual Decision Making. Cerebral Cortex 13, 1257–1269, doi:10.1093/cercor/bhg097 (2003).

14 Lo, C.-C. & Wang, X.-J. Cortico-basal ganglia circuit mechanism for a decision threshold in reaction time tasks. Nat Neurosci 9, 956–963, doi:http://www.nature.com/neuro/journal/v9/n7/suppinfo/nn1722_S1.html (2006).

15 Usher, M. & McClelland, J. L. The time course of perceptual choice: The leaky, competing accumulator model. Psychological Review 108, 550–592, doi:10.1037/0033-295X.108.3.550 (2001).

16 Cisek, P., Puskas, G. A. & El-Murr, S. Decisions in Changing Conditions: The Urgency-Gating Model. The Journal of Neuroscience 29, 11560 (2009).

17 Huk, A. C. & Shadlen, M. N. Neural Activity in Macaque Parietal Cortex Reflects Temporal Integration of Visual Motion Signals during Perceptual Decision Making. The Journal of Neuroscience 25, 10420 (2005).

18 Kiani, R. & Shadlen, M. N. Representation of confidence associated with a decision by neurons in the parietal cortex. Science (New York, N.Y.) 324, 759–764, doi:10.1126/science.1169405 (2009).

19 Smith, P. L. & Ratcliff, R. Psychology and neurobiology of simple decisions. Trends in Neurosciences 27, 161–168, doi:10.1016/j.tins.2004.01.006 (2004).

20 Ditterich, J. Evidence for time-variant decision making. European Journal of Neuroscience 24, 3628–3641, doi:10.1111/j.1460-9568.2006.05221.x (2006).

21 Hanks, T., Kiani, R. & Shadlen, M. N. A neural mechanism of speed-accuracy tradeoff in macaque area LIP. eLife 3, e02260, doi:10.7554/eLife.02260 (2014).

22 Wong, K.-F., Huk, A., Shadlen, M. & Wang, X.-J. Neural circuit dynamics underlying accumulation of time-varying evidence during perceptual decision making. Frontiers in Computational Neuroscience 1, 6 (2007).

23 Inagaki, H. K., Fontolan, L., Romani, S. & Svoboda, K. Discrete attractor dynamics underlies persistent activity in the frontal cortex. Nature 566, 212–217, doi:10.1038/s41586-019-0919-7 (2019).

24 Seidemann, E., Meilijson, I., Abeles, M., Bergman, H. & Vaadia, E. Simultaneously recorded single units in the frontal cortex go through sequences of discrete and stable states in monkeys performing a delayed localization task. The Journal of Neuroscience 16, 752 (1996).

25 Golub, M. D., Chase, S. M., Batista, A. P. & Yu, B. M. Brain–computer interfaces for dissecting cognitive processes underlying sensorimotor control. Current Opinion in Neurobiology 37, 53–58, doi:https://doi.org/10.1016/j.conb.2015.12.005 (2016).

26 Sussillo, D., Stavisky, S. D., Kao, J. C., Ryu, S. I. & Shenoy, K. V. Making brain– machine interfaces robust to future neural variability. Nature Communications 7, 13749, doi:10.1038/ncomms13749 https://www.nature.com/articles/ncomms13749#supplementary-information (2016).

27 Semedo, J. D., Zandvakili, A., Machens, C. K., Yu, B. M. & Kohn, A. Cortical Areas Interact through a Communication Subspace. Neuron 102, 249–259.e244, doi:https://doi.org/10.1016/j.neuron.2019.01.026 (2019).

28 Seideman, J. A., Stanford, T. R. & Salinas, E. Saccade metrics reflect decision-making dynamics during urgent choices. Nature Communications 9, 2907, doi:10.1038/s41467-018-05319-w (2018).

29 Musall, S., Kaufman, M. T., Juavinett, A. L., Gluf, S. & Churchland, A. K. Single-trial neural dynamics are dominated by richly varied movements. bioRxiv, 308288, doi:10.1101/308288 (2019).

30 Aflalo, T. et al. Decoding motor imagery from the posterior parietal cortex of a tetraplegic human. Science 348, 906 (2015).

31 Andersen, R. A., Hwang, E. J. & Mulliken, G. H. Cognitive Neural Prosthetics. Annual Review of Psychology 61, 169–190, doi:10.1146/annurev.psych.093008.100503 (2009).

32 Musallam, S., Corneil, B. D., Greger, B., Scherberger, H. & Andersen, R. A. Cognitive Control Signals for Neural Prosthetics. Science 305, 258 (2004).

33 Pesaran, B., Musallam, S. & Andersen, R. A. Cognitive neural prosthetics. Current Biology 16, R77–R80, doi:10.1016/j.cub.2006.01.043 (2006).

34 Benjamini, Y. & Hochberg, Y. Controlling The False Discovery Rate - A Practical And Powerful Approach To Multiple Testing. Vol. 57 (1995).

