## Supplementary Information for "Decoding and perturbing decision states in real time"

| Linear regression on ΔDV slope | | | | | | | |
|---|---|---|---|---|---|---|---|
| | | Monkey H | | | Monkey F | | |
| | Predictor | Beta value | SE | p-value | Beta value | SE | p-value |
| DV boundary = 1 | Bias | -0.0022 | 0.0036 | 0.5331 | -0.0147 | 0.0030 | 9.38E-07 |
| | Signed coherence | 0.0084 | 0.0005 | 5.20E-70 | 0.0033 | 0.0002 | 3.87E-55 |
| | Pulse onset time | -0.0257 | 0.0146 | 0.0783 | 0.1038 | 0.0188 | 3.21E-08 |
| | Pulse direction | 0.0203 | 0.0036 | 1.27E-08 | 0.0120 | 0.0030 | 5.93E-05 |
| | Pulse onset time × direction | -0.0393 | 0.0146 | 0.0071 | -0.0362 | 0.0188 | 0.0535 |
| DV boundary = 2 | Bias | -0.0224 | 0.0061 | 0.0003 | -0.0033 | 0.0031 | 0.2865 |
| | Signed coherence | 0.0017 | 0.0005 | 0.0006 | 0.0018 | 0.0002 | 4.63E-18 |
| | Pulse onset time | 0.0676 | 0.0143 | 2.46E-06 | 0.0404 | 0.0103 | 9.25E-05 |
| | Pulse direction | 0.0167 | 0.0061 | 0.0066 | 0.0040 | 0.0031 | 0.1915 |
| | Pulse onset time × direction | -0.0286 | 0.0143 | 0.0459 | -0.0138 | 0.0103 | 0.1806 |
| DV boundary = 3 | Bias | 0.0241 | 0.0070 | 0.0006 | 0.0152 | 0.0039 | 8.80E-05 |
| | Signed coherence | -0.0004 | 0.0005 | 0.3928 | 0.0009 | 0.0002 | 1.78E-06 |
| | Pulse onset time | -0.0221 | 0.0136 | 0.1040 | -0.0346 | 0.0092 | 0.0002 |
| | Pulse direction | -0.0040 | 0.0070 | 0.5685 | 0.0023 | 0.0039 | 0.5504 |
| | Pulse onset time × direction | 0.0085 | 0.0136 | 0.5313 | -0.0027 | 0.0092 | 0.7726 |
| DV boundary = 4 | Bias | 0.0347 | 0.0082 | 2.40E-05 | 0.0052 | 0.0052 | 0.3241 |
| | Signed coherence | -0.0012 | 0.0005 | 0.0232 | 0.0007 | 0.0002 | 0.0038 |
| | Pulse onset time | -0.0418 | 0.0144 | 0.0036 | -0.0210 | 0.0104 | 0.0432 |
| | Pulse direction | -0.0002 | 0.0082 | 0.9758 | 0.0093 | 0.0052 | 0.0741 |
| | Pulse onset time × direction | 0.0013 | 0.0144 | 0.9270 | -0.0146 | 0.0104 | 0.1596 |

**Supplementary Information Table 1– Coefficients obtained from linear regression on ΔDV slope, motion pulse perturbation experiment (monkeys H and F)**

| Logistic regression on choice | | | | | | | |
|---|---|---|---|---|---|---|---|
| | | Monkey H | | | Monkey F | | |
| | **Predictor** | **Beta value** | **SE** | **p-value** | **Beta value** | **SE** | **p-value** |
| **DV boundary = 1** | Bias | 0.3247 | 0.0587 | 3.19E-08 | -0.2165 | 0.0549 | 7.98E-05 |
| | Signed coherence | 0.2623 | 0.0091 | 4.73E-184 | 0.0986 | 0.0041 | 4.68E-127 |
| | Pulse onset time | -1.1533 | 0.2391 | 1.41E-06 | -0.9315 | 0.3468 | 0.0072 |
| | Pulse direction | 0.4844 | 0.0588 | 1.80E-16 | 0.2552 | 0.0549 | 3.30E-06 |
| | Pulse onset time × direction | -1.0683 | 0.2389 | 7.79E-06 | -0.9993 | 0.3467 | 0.0040 |
| **DV boundary = 2** | Bias | 0.2599 | 0.1048 | 0.0131 | -0.4291 | 0.0587 | 2.62E-13 |
| | Signed coherence | 0.3502 | 0.0110 | 3.98E-222 | 0.1031 | 0.0042 | 1.94E-130 |
| | Pulse onset time | -0.8048 | 0.2422 | 0.0009 | 0.5168 | 0.1949 | 0.0080 |
| | Pulse direction | 0.1648 | 0.1048 | 0.1157 | 0.1829 | 0.0586 | 0.0018 |
| | Pulse onset time × direction | -0.2920 | 0.2421 | 0.2277 | -0.4518 | 0.1947 | 0.0203 |
| **DV boundary = 3** | Bias | 0.2083 | 0.1363 | 0.1264 | -0.6655 | 0.0813 | 2.79E-16 |
| | Signed coherence | 0.3399 | 0.0116 | 3.93E-188 | 0.1071 | 0.0045 | 4.47E-123 |
| | Pulse onset time | -0.6986 | 0.2610 | 0.0074 | 0.9975 | 0.1924 | 2.15E-07 |
| | Pulse direction | 0.1669 | 0.1362 | 0.2203 | 0.1036 | 0.0811 | 0.2015 |
| | Pulse onset time × direction | -0.1851 | 0.2607 | 0.4777 | -0.2748 | 0.1922 | 0.1529 |
| **DV boundary = 4** | Bias | -0.0200 | 0.1783 | 0.9106 | -0.6476 | 0.1101 | 4.07E-09 |
| | Signed coherence | 0.3834 | 0.0154 | 1.12E-136 | 0.1028 | 0.0053 | 7.34E-84 |
| | Pulse onset time | -0.4523 | 0.3109 | 0.1457 | 1.3977 | 0.2198 | 2.04E-10 |
| | Pulse direction | 0.0512 | 0.1784 | 0.7742 | 0.1174 | 0.1100 | 0.2855 |
| | Pulse onset time × direction | -0.0181 | 0.3109 | 0.9536 | -0.1738 | 0.2197 | 0.4289 |

**Supplementary Information Table 2– Coefficients obtained from logistic regression on choice, motion pulse perturbation experiment (monkeys H and F)**

| Subject | Session | 1 | 2 | 3 | 4 | 5 | 6 | 7 | 8 | 9 | 10 | 11 | 12 | 13 | 14 | 15 | 16 | 17 |
|---|---|---|---|---|---|---|---|---|---|---|---|---|---|---|---|---|---|---|
| **Monkey H** | tminPre(s) | 0.1 | 0.1 | 0.1 | 0.1 | 0.05 | 0.05 | 0.05 | 0.05 | 0.05 | 0.05 | 0.15 | 0.15 | 0.15 | 0.1 | 0.05 | 0.15 | 0.15 |
| | tminPost(s) | 0.1 | 0.1 | 0.1 | 0.1 | 0.05 | 0.05 | 0.05 | 0.05 | 0.05 | 0.05 | 0.15 | 0.15 | 0.15 | 0.15 | 0.15 | 0.05 | 0.1 |
| | DVminPre | 2 | 1.5 | 3 | 1 | 2 | 1 | 1.5 | 3 | 1.5 | 1 | 2 | 2 | 2 | 2 | 2 | 2 | 2 |
| | DVminPost | 2 | 1.5 | 3 | 1 | 2 | 1 | 1.5 | 3 | 0.1 | 0.1 | 2 | 2 | 2 | 2 | 2 | 2 | 2 |
| | tminDots(s) | 0.25 | 0.25 | 0.25 | 0.25 | 0.25 | 0.25 | 0.25 | 0.25 | 0.25 | 0.25 | 0.25 | 0.25 | 0.25 | 0.25 | 0.25 | 0.25 | 0.25 |
| **Monkey F** | tminPre(s) | 0.1 | 0.1 | 0.1 | 0.1 | 0.1 | 0.1 | 0.05 | 0.05 | 0.05 | 0.05 | 0.05 | 0.05 | 0.05 | 0.15 | 0.15 | | |
| | tminPost(s) | 0.1 | 0.1 | 0.1 | 0.1 | 0.1 | 0.1 | 0.05 | 0.05 | 0.05 | 0.05 | 0.05 | 0.05 | 0.05 | 0.05 | 0.1 | | |
| | DVminPre | 2 | 2 | 2 | 1.5 | 3 | 1 | 2 | 3 | 1.5 | 1 | 1 | 1.5 | 1 | 2 | 2 | | |
| | DVminPost | 2 | 2 | 2 | 1.5 | 3 | 1 | 2 | 3 | 1.5 | 1 | 1 | 0.1 | 0.1 | 2 | 2 | | |
| | tminDots(s) | 0.25 | 0.25 | 0.25 | 0.25 | 0.25 | 0.25 | 0.25 | 0.25 | 0.25 | 0.25 | 0.25 | 0.25 | 0.25 | 0.25 | 0.25 | | |

**Supplementary Information Table 3– Parameters used for virtual boundaries and CoM closed loop experiments (monkeys H and F)**
